# Protease-Activated Receptor 1 as an Endogenous Model of Peptidergic Gαq-Gα12-Biased G Protein Signaling

**DOI:** 10.64898/2026.03.03.709348

**Authors:** Braden S. Fallon, Robert A. Campbell, Justin G. English

## Abstract

G protein-coupled receptors (GPCRs) are the most widely targeted class of signaling proteins, comprising ~30% of FDA-approved drugs. Their therapeutic potential arises from their ability to translate diverse extracellular cues into intracellular signals via G proteins, arrestins, and other effectors. This signaling versatility creates opportunities for functional selectivity, where ligands preferentially engage particular pathways. However, few endogenous receptor systems display well-defined ligand-dependent divergence across multiple signaling levels. Protease-activated receptor 1 (PAR1) is a candidate model. Thrombin canonically cleaves PAR1 at Arg41, whereas activated protein C (aPC) has been reported to cleave PAR1 at Arg46 in endothelial and co-receptor-supported settings, creating distinct tethered peptide ligands. Thrombin cleavage drives canonical Gαq- and Gα12/13-dependent prothrombotic and barrier-disruptive signaling, whereas aPC cleavage has been associated with anticoagulant, cytoprotective, and anti-inflammatory signaling. However, PAR1’s transducer-wide coupling profile, transcriptional consequences, and physiological outputs remain incompletely characterized. We integrated transducer-wide biosensor assays (TRUPATH, TGFα shedding, PRESTO-Tango), analysis of a PAR1 ± thrombin TRE-MPRA dataset followed by targeted TRE dual-luciferase validation, and platelet activation and calcium flux studies in primary human platelets to define how protease identity reshapes signaling from proximal transducer engagement to physiological output. Thrombin produced robust PAR1 coupling to Gαq and Gα12, whereas aPC produced detectable coupling only to Gα12. Neither protease generated detectable β-arrestin-2 recruitment in PRESTO-Tango. Both proteases supported dose-dependent TGFα shedding that was insensitive to FR900359. A PAR1 ± thrombin TRE-MPRA dataset identified thrombin-responsive transcriptional response elements and nominated NFκB1 and THRB for targeted follow-up. Luciferase assays showed NFκB1 was thrombin-induced and FR900359-sensitive, whereas THRB was induced by both thrombin and aPC and was FR900359-insensitive. In primary human platelets, thrombin, but not aPC, induced P-selectin expression and a calcium response. Thrombin responses were suppressed by FR900359, supporting a requirement for Gαq in these platelet activation markers. Together, these findings support PAR1 as an endogenous model of protease-dependent functional selectivity that, in our heterologous assay systems, separates signaling along a Gαq-versus-Gα12 axis, thus providing a framework for future technologies, such as high-throughput tethered-peptide evolution platforms, to understand the principles of G protein selectivity across GPCRs.

## INTRODUCTION

G protein-coupled receptors (GPCRs) are the most extensively targeted family of signaling proteins in modern pharmacology, with ~30% of FDA-approved drugs acting through these receptors (Hauser et al., 2017; Hilger et al., 2018; Casadó and Casadó-Anguera, 2023; Zhang et al., 2024; Lorente et al., 2025). The therapeutic promise of GPCRs lies not only in their ubiquity across physiological systems but also in their ability to transduce diverse extracellular cues into intracellular signals through G proteins, arrestins, and other effectors, thereby regulating processes including cardiovascular function, neurotransmission, and immune signaling (Hauser et al., 2017; Besserer-Offroy et al., 2018). This signaling versatility provides opportunities for functional selectivity (biased agonism), in which distinct ligands stabilize receptor conformations that preferentially activate one signaling pathway over another (St-Pierre et al., 2018).

Over the past two decades, functional selectivity has emerged as a leading strategy for developing safer and more effective GPCR therapeutics (Hauser et al., 2017). Several biased agonists have entered clinical development—including TRV130/oliceridine, a μ-opioid receptor ligand designed to reduce β-arrestin recruitment while preserving G protein signaling (Chen et al., 2013; DeWire et al., 2013), and TRV027, an angiotensin II type 1 receptor ligand designed to bias away from Gαq coupling (Violin et al., 2010). Despite promising preclinical data, both compounds failed to demonstrate clear advantages in clinical trials. Oliceridine showed dose-limiting respiratory depression similar to that of morphine (Singla et al., 2019), and TRV027 did not improve outcomes in patients with acute heart failure (Pang et al., 2017). More broadly, multiple dopamine and serotonin receptor-biased ligands have struggled to achieve clinical endpoints despite compelling mechanistic rationale (Michel and Charlton, 2018; Newman-Tancredi et al., 2022). These outcomes highlight a central challenge: the field lacks well-characterized endogenous GPCR systems that naturally exhibit bias and can serve as model platforms for translational discovery.

Protease-activated receptor 1 (PAR1) represents a compelling candidate to fill this gap. It is an endogenous GPCR with multiple native protease ligands that can generate discrete, protease-defined receptor states. PAR1 plays a central role in hemostasis as the primary thrombin receptor on human platelets (Vu et al., 1991; Coughlin, 2005). Its unusual activation mechanism—proteolytic cleavage of the N-terminus to reveal a tethered agonist— creates an intrinsic opportunity for functional selectivity, as cleavage at different positions can expose distinct tethered peptides and thereby specify different signaling programs at the same receptor (Cottrell et al., 2002; Soh and Trejo, 2011). Thrombin cleaves PAR1 at Arg41 to reveal a tethered ligand that canonically drives Gαq- and Gα12/13-dependent signaling, promoting platelet aggregation, endothelial permeability, and vascular remodeling (Hung et al., 1992; Offermanns et al., 1994; McCoy et al., 2010; Guo et al., 2024). In contrast, activated protein C (aPC) has been reported to cleave PAR1 at the noncanonical Arg46 site in endothelial and co-receptor-supported settings, where cytoprotective and anti-inflammatory outputs have been observed, often in the presence of endothelial protein C receptor (EPCR) and thrombomodulin (Ludeman et al., 2005; Mosnier et al., 2012; Rezaie, 2014; De Ceunynck et al., 2018; Stojanovski and Di Cera, 2025). Importantly, because PAR1 is expressed in both platelets and endothelial cells, thrombin and aPC can be compared within each cellular context. In platelets, thrombin elicits strong PAR1-dependent activation phenotypes. In the endothelium, thrombin and aPC likewise drive divergent barrier and inflammatory outcomes. This protease-dependent duality provides a biologically relevant model for functional selectivity and a direct link to cardiovascular disease, thrombosis, and inflammation—key areas of unmet clinical need (Rezaie, 2014; Antoniak et al., 2021). Consistent with this, PAR1-biased signaling has been implicated in endothelial dysfunction in sickle cell disease, where distinct PAR1 agonist profiles modulate vascular inflammation and barrier function (Ramadas et al., 2024).

Despite its promise, the signaling landscape of PAR1 remains incompletely mapped. Much of the literature has relied on second messenger assays or endpoint functional readouts, which provide only partial insight into pathway selectivity and transducer coupling (van den Eshof et al., 2017; Han et al., 2020; Willis Fox and Preston, 2020). More recently, effector membrane translocation (EMTA) biosensors and structure-based GPCR coupling atlases have begun to survey G protein and β-arrestin coupling profiles for large receptor panels, including PAR1 (Avet et al., 2022; Hauser et al., 2022; Pándy-Szekeres et al., 2024). However, these approaches do not typically connect transducer-wide coupling to downstream transcriptional programs or physiological outputs, nor do they explicitly compare endogenous protease ligands at a single receptor. Comprehensive analyses that track information flow from G protein coupling to gene expression and cell-level phenotypes are needed to establish PAR1 as a G protein-selective model and to clarify the mechanistic basis of its divergent biological effects.

Here, we address this gap using a multi-platform strategy that follows PAR1 from proximal coupling events to integrated cellular phenotypes. We combine direct transducer activation assays—including TRUPATH for subtype-resolved G protein coupling (Olsen et al., 2020), TGFα shedding for proximal Gαq/Gα12 responses (Inoue et al., 2012, 2019), and PRESTO-Tango for β-arrestin recruitment (Kroeze et al., 2015)—with analysis of a PAR1 ± thrombin transcriptional response element massively parallel reporter dataset (Zahm et al., 2024) and platelet activation measurements. This framework enables pathway-level mapping of thrombin- and aPC-mediated PAR1 signaling across recombinant HEK/HTLA biosensor systems and primary human platelets.

We find that thrombin engages both Gαq and Gα12, whereas aPC produces detectable coupling only to Gα12 under the same assay conditions. Under these experimental conditions, neither protease induces detectable β-arrestin-2 recruitment by PRESTO-Tango. These divergent coupling profiles align with distinct transcriptional reporter signatures and platelet activation phenotypes. Together, our results support PAR1 as a useful endogenous model of protease-dependent functional selectivity and provide a framework for future efforts— such as high-throughput ligand screening—to engineer ligands with defined positions in G protein coupling space.

## RESULTS

### Thrombin Drives Canonical Gαq/Gα12 Signaling Through PAR1

PAR1 has historically been described as a Gαq- and Gα12/13-coupled receptor in the context of thrombin signaling, but these conclusions have largely arisen from pathway-specific readouts rather than direct, systematic profiling of individual Gα subunits (Hung et al., 1992; Offermanns et al., 1994; McCoy et al., 2010; Willis Fox and Preston, 2020). To comprehensively define the coupling landscape of thrombin-activated PAR1, we first validated the TRUPATH platform in HEK293 cells transiently expressing the indicated receptor constructs. TRUPATH is a BRET-based biosensor system that enables quantitative, subtype-resolved measurement of GPCR-Gα interactions in live cells (Olsen et al., 2020). Using three key GPCRs— neurotensin receptor 1 (NTSR1), kappa-opioid receptor (OPRK), and beta-2 adrenergic receptor (ADRB2)—we benchmarked all sensor constructs based on their specific Gα coupling preferences. As expected, NTSR1 displayed robust coupling across multiple Gα (Gαi1, Gαi2, Gαi3, GαoA, GαoB, Gαz, Gαq, Gα11, Gα12, Gα13, and Gα15). Gαs-short coupling was confirmed with ADRB2 and OPRK demonstrated strong engagement of Gαgustducin, collectively verifying assay performance (**Figure 1A, Supplementary Figure S1, Supplementary Table S1**).

**Figure 1.**
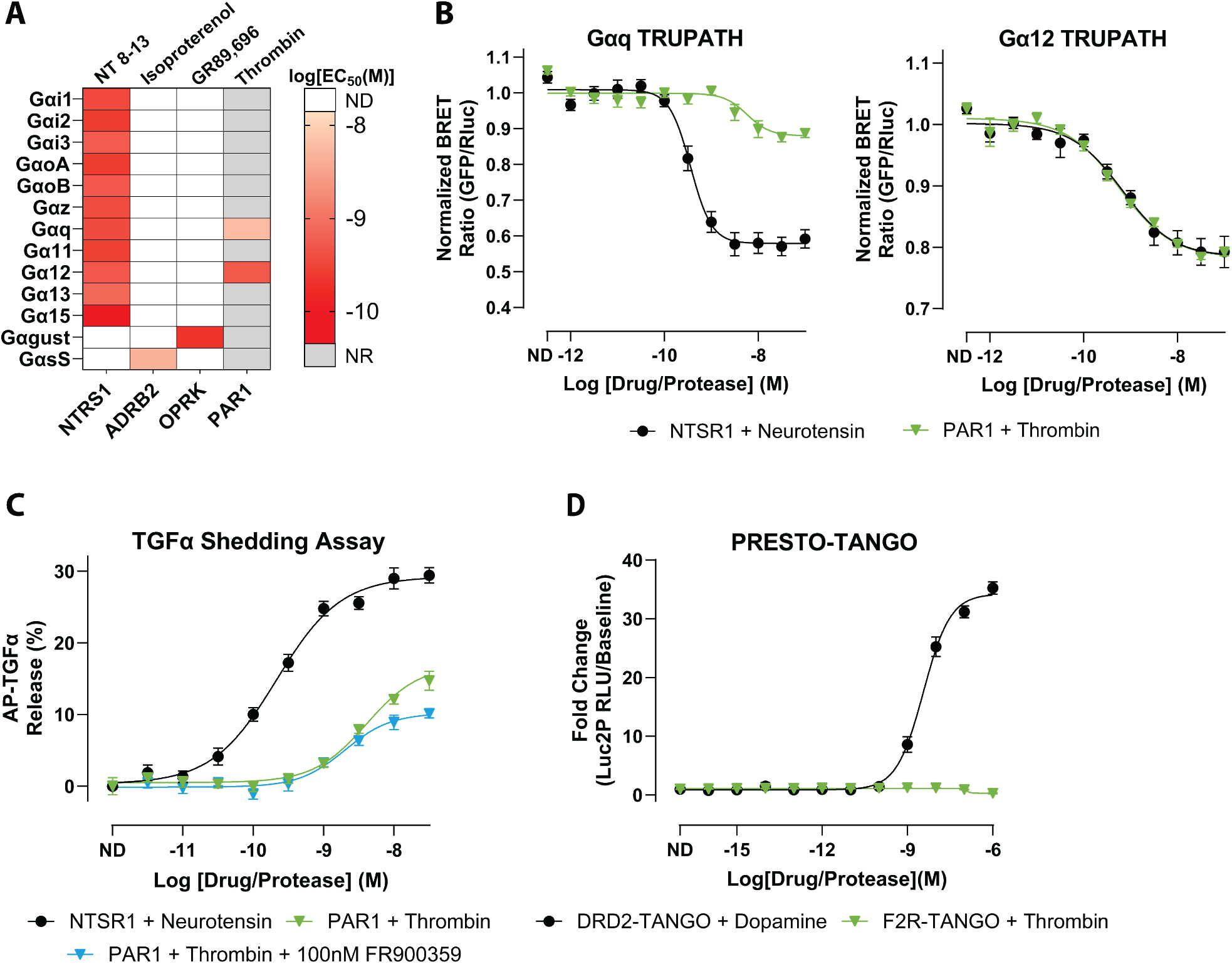
Thrombin-activated PAR1 signals through Gαq and Gα12. **A)** Heatmap of EC_50_ values for the 13 Gα subunits determined from TRUPATH concentration-response fits. The map displays EC_50_ values for the positive-control receptors included in the study, along with PAR1 responses with thrombin. Each cell corresponds to the EC_50_ (M) derived from the concentration-response curve for the indicated Gα. Color intensity denotes potency. Grey boxes indicate Gα/receptor pairs for which no response was observed (NR). White boxes denote pairs for which coupling was not determined (ND). **B)** Representative TRUPATH concentration-response curves for Gαq (left) and Gα12 (right). Curves compare NTSR1 stimulation with neurotensin and PAR1 stimulation with thrombin. Symbols show mean ± SEM across biological replicates; curves are nonlinear fits to the pooled replicate data used to extract EC_50_ values. **C)** TGFα-shedding assay percentage of released AP-TGFα normalized to baseline, showing dose-dependent responses of PAR1 with thrombin in the presence or absence of FR900359. NTSR1 with neurotensin serves as the positive control. Data are shown as the mean signal ± SEM. **D)** PRESTO-Tango β-Arrestin-2 recruitment assay comparing fold change of DRD2 stimulation with dopamine and PAR1 stimulation with thrombin. Data are plotted as mean ± SEM. All plotted points represent the mean of three independent biological replicates, each measured in quadruplicate (N = 3, n = 4).

These data were consistent with previously reported coupling patterns and potencies for prototypic GPCR controls and with prior transducer-wide GPCR–G protein coupling resources (Olsen et al., 2020; Avet et al., 2022; Hauser et al., 2022; Pándy-Szekeres et al., 2024).

To confirm that our TRUPATH acquisition window captured enzymatic activation of PAR1 by thrombin, we performed a time-course experiment monitoring PAR1-Gα12 BRET after thrombin addition (**Supplementary Figure S2**). The BRET signal stabilized after 10 minutes; thus, no modification to the TRUPATH protocol was necessary as the enzymatic activity would fall within our standard kinetic acquisition window. Across all 13 TRUPATH sensors, thrombin-activated PAR1 produced robust coupling to Gαq and Gα12, with no detectable activation of other Gα families within the sensitivity of our assay (**Figure 1A-B, Supplementary Figure S1, Supplementary Table S1**). Concentration-response analysis revealed that thrombin displayed higher apparent potency for Gα12 than for Gαq (LogEC_50_ = −9.29 vs −8.24) (**Figure 1B**). This left shift suggests that near-maximal Gα12 engagement was reached at lower thrombin concentrations than near-maximal Gαq engagement. Trends were observed for some non-primary sensors, including Gα13, but these responses were smaller and less reproducible than those of Gαq and Gα12 and did not support robust concentration-response fitting in this dataset. BRET responses for all other Gα subunits remained at baseline (**Supplementary Figure S1**). Together, these data suggest that in HEK293 cells, thrombin-activated PAR1 is functionally restricted to Gαq and Gα12.

As an orthogonal validation for Gα12 signaling, we employed the TGFα shedding assay, a metalloprotease-dependent ectodomain release reporter of GPCR activation that integrates Gα12/13-mediated signaling (Inoue et al., 2012, 2019). HEK293 cells transiently expressed AP-TGFα and the receptor constructs. Thrombin stimulation of PAR1 produced a clear dose-dependent increase in shedding (LogEC_50_ = −8.39 ± 0.10, Span = 16.9 ± 1.46%) (**Figure 1C, Supplementary Table S2**).

To test whether the observed effector responses required Gαq-family signaling, we used FR900359, a selective inhibitor of Gαq/11/14 (Schrage et al., 2015; Reher et al., 2018). In a representative TRUPATH control experiment using NTSR1, FR900359 abolished detectable Gαq/11 activation while preserving signaling through non-Gαq sensors (**Supplementary Figure S3**), consistent with prior reports.

We found that with the addition of FR900359 to the assay, thrombin stimulation of PAR1 exhibited a similar potency to the non-FR900359-treated condition and showed a minor decrease in response amplitude (LogEC_50_ = −8.69 ± 0.11, Span = 10.2 ± 0.85%) (**Figure 1C, Supplementary Table S2**).

We next examined β-arrestin-2 recruitment using PRESTO-Tango, which reports β-arrestin engagement through a TEV protease-dependent transcriptional readout in HTLA cells expressing F2R-Tango or DRD2-Tango constructs. In contrast to the robust dopamine-induced signal observed with the DRD2 positive control, thrombin stimulation of PAR1 produced no measurable increase in β-arrestin-2-dependent reporter activity across the tested concentration range (**Figure 1D, Supplementary Table S3**). Thus, in this assay format and cell context, thrombin-activated PAR1 exhibits strong G protein coupling (Gαq/Gα12) without detectable β-arrestin-2 recruitment.

Because TRAP-6 is a soluble peptide agonist that mimics the canonical thrombin-revealed PAR1 tethered ligand while bypassing proteolytic cleavage, we tested TRAP-6 in TRUPATH and PRESTO-Tango assays as a non-proteolytic comparator of thrombin-like PAR1 activation. In TRUPATH, TRAP-6 produced detectable PAR1 coupling to both Gαq and Gα12, with higher apparent potency at Gα12 (LogEC_50_ = −6.37 ± 0.11; EC50 = 0.43 µM) than at Gαq (LogEC_50_ = −5.28 ± 0.16; EC50 = 5.29 µM). In PRESTO-Tango, TRAP-6 did not produce a detectable β-arrestin-2 recruitment signal under the assay conditions tested (**Supplementary Figure S4, Supplementary Table S4**). These TRAP-6 data provide a receptor-intrinsic, non-proteolytic comparator showing that under the assay conditions used, PAR1 can produce detectable Gαq and Gα12 coupling without a detectable PRESTO-Tango β-arrestin-2 signal.

Collectively, these data indicate that thrombin-activated PAR1 is strongly G protein-biased in HEK293/HTLA cells, with detectable coupling limited to Gαq and Gα12 and no detectable β-arrestin-2 recruitment by PRESTO-Tango under these conditions.

### aPC Restricts PAR1 Signaling to Gα12 and Bypasses Gαq Activation

We next applied the same TRUPATH profiling to PAR1 in the presence of activated protein C (aPC) to determine how protease identity reshapes the receptor’s coupling profile. In contrast to thrombin, aPC produced detectable coupling of PAR1 only to Gα12, with no measurable activation of Gαq or other Gα subtypes across the tested concentrations (**Figure 2A-B, Supplementary Figure S1, Supplementary Table S1**). The potency of aPC for Gα12 was right-shifted relative to thrombin (LogEC_50_ = −8.61), indicating weaker apparent agonism while remaining clearly active. Responses for all other TRUPATH sensors were indistinguishable from baseline, consistent with a Gα12-biased PAR1 signaling state under these assay conditions.

**Figure 2.**
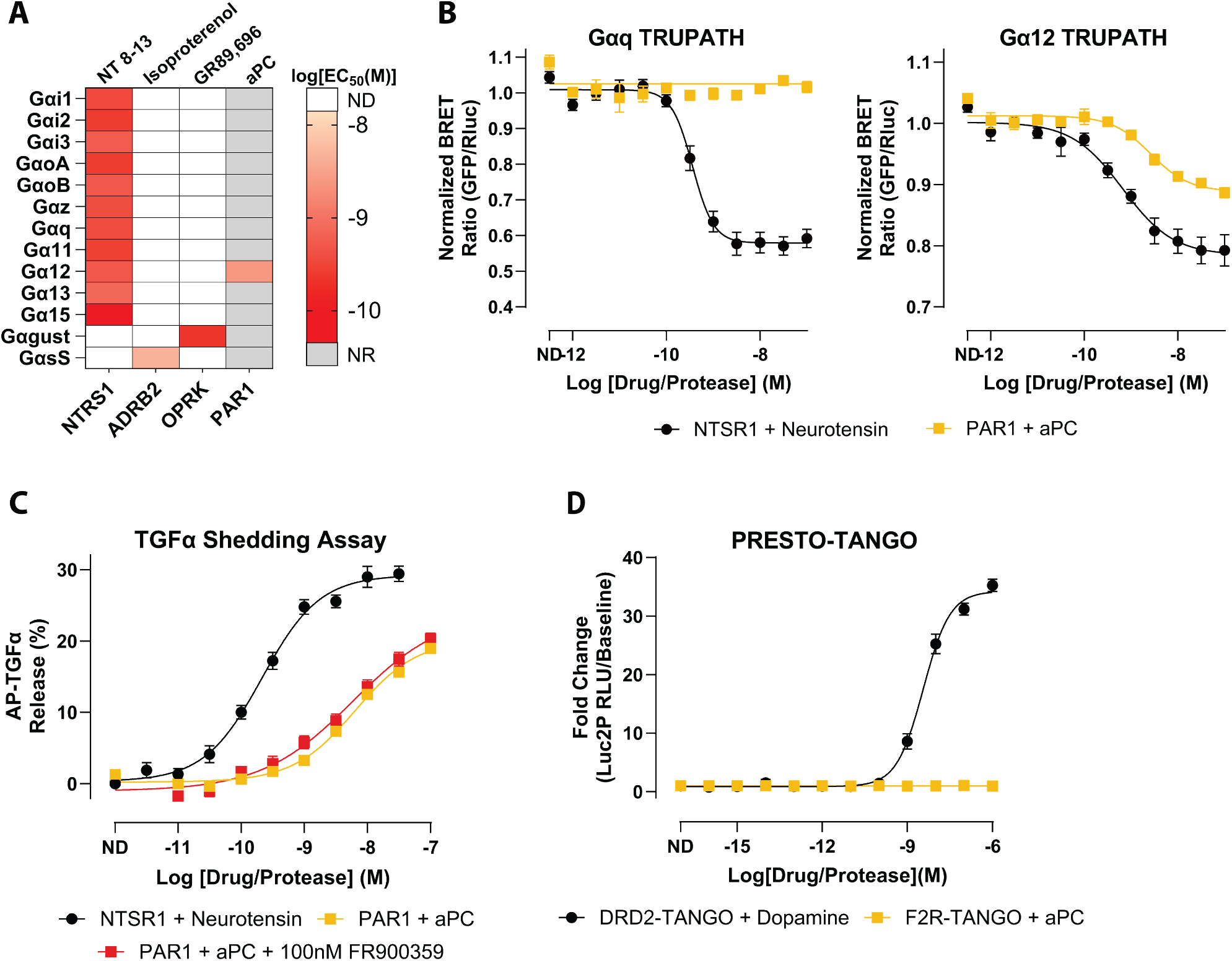
Activated protein C (aPC) exhibits PAR1 coupling to only Gα12. **A)** Heatmap of EC_50_ values for the 13 individual Gα subunits determined from TRUPATH concentration-response fits. The map displays EC_50_ values for the positive-control receptors included in the study, along with PAR1 responses measured using activated protein C (aPC). Each cell corresponds to the EC_50_ (M) calculated from the concentration-response curve for the indicated Gα. Color intensity denotes potency. Grey boxes indicate Gα/receptor pairs for which no response was observed (NR). White boxes denote pairs for which coupling was not determined (ND). **B)** Representative TRUPATH concentration-response curves for Gαq (left) and Gα12 (right). Curves compare NTSR1 stimulation with neurotensin and PAR1 stimulation with aPC. Symbols show mean ± SEM across biological replicates; curves are nonlinear fits to the pooled replicate data used to extract EC_50_ values. **C)** TGFa-shedding assay percentage of released AP-TGFα normalized to baseline, showing dose-dependent responses of PAR1 with aPC in the presence or absence of FR900359. NTSR1 with neurotensin serves as the positive control. Data are shown as the mean signal ± SEM. **D)** PRESTO-Tango β-Arrestin-2 recruitment assay comparing fold change of DRD2 stimulation with dopamine and PAR1 stimulation with aPC. Data are plotted as mean ± SEM. All plotted points represent the mean of three independent biological replicates, each measured in quadruplicate (N = 3, n = 4).

We then asked whether this Gα12-only profile was sufficient to drive downstream G protein-dependent effector responses. In the TGFα shedding assay, aPC-activated PAR1 produced a robust, dose-dependent response comparable in magnitude to that of thrombin (LogEC_50_ = −8.19 ± 0.07, Span = 20.2 ± 1.03%) (**Figure 2C, Supplementary Table S2**). FR900359 pretreatment followed by aPC stimulation of PAR1 exhibited similar potency and response amplitude to the non-FR900359-treated condition (LogEC_50_ = −8.23 ± 0.19, Span = 24.1 ± 2.82%) (**Figure 2C, Supplementary Table S2**). The similarity in potency and response amplitude between aPC and thrombin in this assay indicates that Gα12 alone is sufficient to support TGFα shedding, and that Gαq engagement is not required for this specific proximal signaling outcome in HEK293 cells.

PRESTO-Tango similarly showed no detectable β-arrestin-2 recruitment in response to aPC across the tested concentrations (**Figure 2D, Supplementary Table S3**), mirroring the arrestin-independent profile observed with thrombin. Thus, in our HEK293/HTLA assay systems, both proteases produced G protein-biased PAR1 signaling, but with distinct patterns of G protein engagement. Thrombin activates both Gαq and Gα12, whereas aPC produces detectable coupling only to Gα12.

Using thrombin as a reference ligand, aPC therefore displays protease-dependent functional selectivity along a Gαq-versus-Gα12 axis. Its relative loss of Gαq coupling while preserving Gα12 activity establishes a naturally occurring Gα12-biased agonist state for PAR1.

### Thrombin-Responsive TREs Nominate Reporters That Distinguish Thrombin Versus aPC Signaling

To connect protease-dependent coupling profiles to downstream transcription, we analyzed a PAR1 ± thrombin TRE-MPRA dataset (Zahm et al., 2024). We use this dataset here as a reporter-discovery step rather than as a stand-alone mechanistic proof. From the extracted data, we focused on the highest-responding promoter architecture (set 1, rotation 8, minCMV). Within this architecture, TREs were ranked by the magnitude of thrombin-induced change (absolute log2 fold change) and statistical support, and the top 10 responses (largest positive and negative changes) were highlighted as candidate thrombin-responsive elements (**Figure 3A-B**).

**Figure 3.**
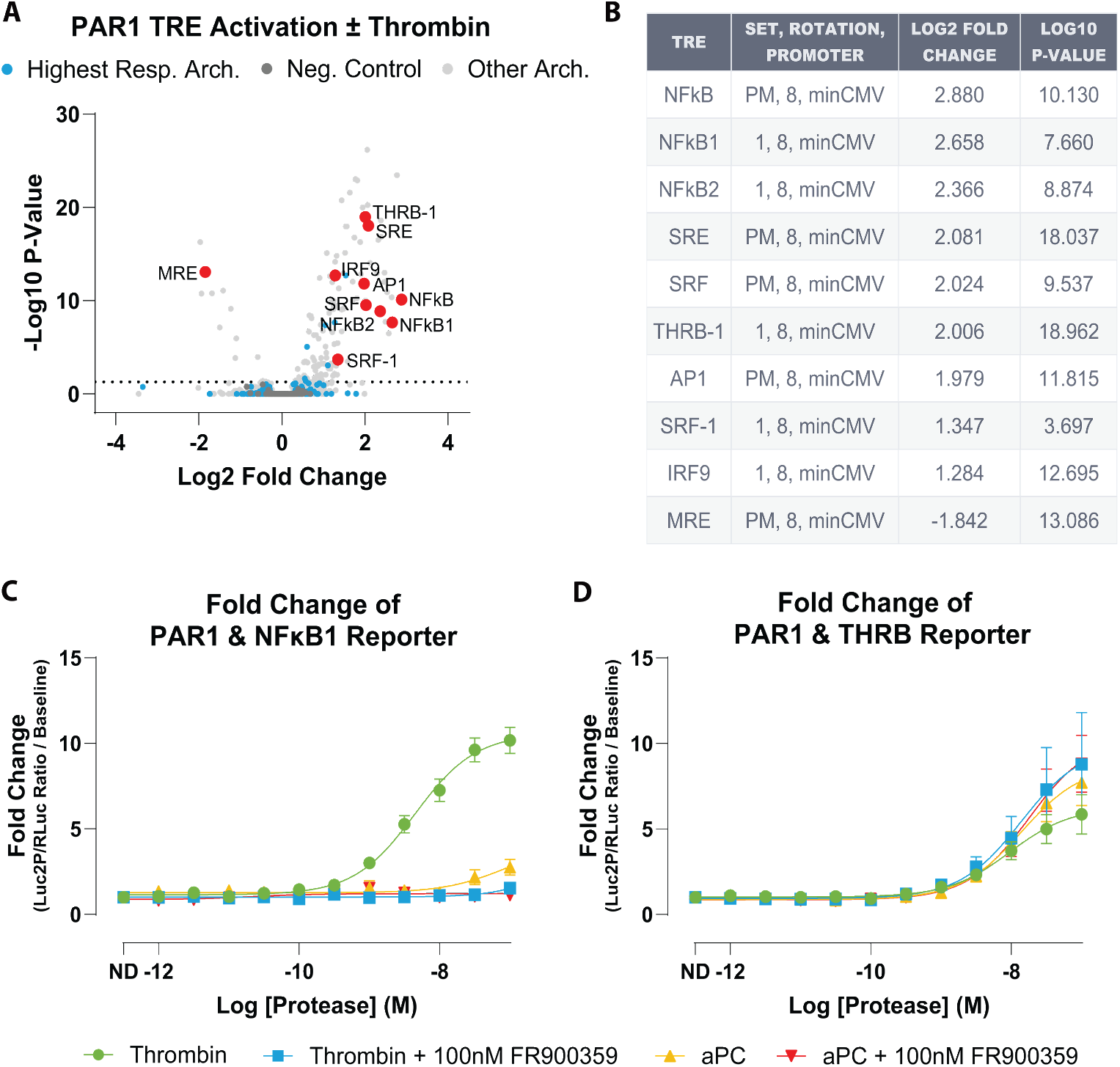
TRE-MPRA dataset nominates PAR1-responsive transcriptional elements, with validation revealing Gαq-dependent NFκB1 and Gαq-independent THRB reporters. **A)** Volcano plot of differential TRE activity in PAR1-expressing HEK293 cells stimulated with thrombin versus vehicle. Each point represents an individual TRE. Light blue points mark TREs belonging to the highest-responding promoter architecture used for downstream analysis, light gray points are TREs from all other architectures, and dark gray points are negative-control TREs. Red points highlight the top 10 thrombin-responsive TREs within the highest-responding architecture. **B)** Summary table of the top 10 TREs, including element identity, promoter architecture, and differential activity statistics. **C)** Individual luciferase validation of the NFκB1 TRE in HEK293 cells expressing PAR1, stimulated with thrombin or activated protein C (aPC) in the presence or absence of the Gαq-family inhibitor FR900359. NFκB1 activity is robustly induced by thrombin, abolished by FR900359, and not detectably induced by aPC. **D)** Individual luciferase validation of the THRB TRE under the same conditions. Both thrombin and aPC induce THRB reporter activity, and THRB is not suppressed by FR900359. Data are plotted as mean ± SEM. All plotted points represent the mean of three independent biological replicates, each measured in quadruplicate (N = 3, n = 4).

From this top 10 set, we selected NFκB1 and THRB as candidate reporters. NFκB1 and THRB refer to synthetic TRE reporter constructs named for the corresponding response-element motifs and are used here as downstream transcriptional readouts of PAR1 signaling, not as direct measurements of endogenous NFκB1 or THRB pathway activity. Importantly, these candidate reporters did not arise solely from the PAR1-specific thrombin dataset. In the broader TRE-MPRA study built from the same synthetic promoter library, individual promoters were orthogonally converted into dual-luciferase reporters (TRE-driven Luc2P firefly with an internal SV40-driven Renilla control) and shown to produce dose-dependent transcriptional responses across multiple stimuli and receptor contexts. Those validation experiments also demonstrated that reporter output depends on promoter architecture, including the paired minimal promoter, indicating that TRE identity and construct context jointly shape signal detection.

Within that broader framework, NFκB1 and THRB emerged as especially informative GPCR-responsive elements. NFκB1 behaved as an FR900359-sensitive readout downstream of Gαq-coupled receptor activation. In contrast, THRB could remain active in receptor contexts where Gαq inhibition suppressed NFκB1, suggesting that these reporters capture separable downstream signaling programs. In addition, the THRB motif was noted to closely resemble an SRF/CArG-like element, raising the possibility that this reporter reflects SRF/RhoA-linked transcriptional output rather than canonical THRB nuclear receptor signaling (Sun et al., 2006; Zahm et al., 2024). Taken together, these prior observations provided a mechanistic rationale for prioritizing NFκB1 and THRB for PAR1 follow-up and for testing whether thrombin and aPC would differentially engage Gαq-sensitive versus Gαq-insensitive transcriptional branches.

Consistent with these observations and our TRUPATH coupling profiles, we predicted that NFκB1 would function as a thrombin-induced, FR900359-sensitive reporter, whereas THRB would report an FR900359-insensitive branch that could be engaged by both thrombin and aPC (Rahman et al., 2002; Vogt et al., 2003; Jin et al., 2009; Gambaryan et al., 2010; Yu and Brown, 2015). To test this, we cotransfected HEK293 cells with PAR1 and the previously generated NFκB1 or THRB dual-luciferase reporter constructs, then stimulated cells with thrombin or aPC in the presence or absence of FR900359.

In this targeted assay, thrombin produced robust NFκB1 reporter activation (LogEC_50_ = −8.32 ± 0.09; Span = 9.79 ± 0.71-fold over vehicle), whereas aPC did not measurably stimulate NFκB1. Pretreatment with FR900359 abolished the thrombin response, consistent with NFκB1 activation requiring Gαq-family signaling under these conditions (**Figure 3C, Supplementary Table S5**). In contrast, the THRB reporter was induced by both thrombin and aPC and was insensitive to FR900359. Thrombin and aPC showed similar potencies through THRB under control conditions (LogEC_50_ = −8.02 ± 0.17 and −7.86 ± 0.15, respectively) and after FR900359 (LogEC_50_ = −7.86 ± 0.28 and −7.75 ± 0.15), with response amplitudes ranging from ~5-10 fold across conditions (**Figure 3D, Supplementary Table S5**). Notably, maximal THRB responses were maintained—and modestly increased—after FR900359 P-(**Supplementary Table S5**).

These data recapitulate the behavior of NFκB1 and THRB reported by Zahm et al., with NFκB1 acting as a Gαq-dependent reporter and THRB reflecting an alternative, FR900359-insensitive signaling branch (Zahm et al., 2024). Given that aPC produces detectable coupling only to Gα12 in TRUPATH, THRB activation is consistent with a Gα12-associated branch in this system.

Thus, analysis of the PAR1 ± thrombin TRE-MPRA dataset, followed by targeted individual TRE validation with thrombin and aPC, supports at least two separable transcriptional reporter programs downstream of PAR1. NFκB1 behaved as a thrombin-induced, FR900359-sensitive reporter, whereas THRB was induced by both thrombin and aPC and was not suppressed by FR900359. These reporter fingerprints mirror the divergence observed at the level of proximal G protein coupling and provide practical surrogates for Gαq-versus Gα12-linked signaling in this system.

### Distinct PAR1 Signaling Modes Produce Divergent Platelet Activation Phenotypes

Given the central role of PAR1 in hemostasis, we next asked how thrombin and aPC signaling modes translate into functional outcomes in primary human platelets. Platelet activation was quantified in five independent human donors by measuring P-selectin surface expression, a well-established marker of α-granule degranulation and platelet activation. Integrin αIIb activation was also assessed as a complementary marker of platelet activation (**Supplementary Figure S5B and S5D**) (Andrianova et al., 2026).

Consistent with canonical PAR1-mediated platelet activation, thrombin induced robust, dose-dependent upregulation of P-selectin in human platelets (**Figure 4A**). To determine the contribution of Gαq signaling, platelets were treated with increasing concentrations of the selective Gαq inhibitor FR900359 (0.5-2.5 µM; **Supplementary Figure S5A, Supplementary Table S6, Supplementary Table S7**). Because 0.5 µM FR900359 produced maximal suppression of thrombin-induced responses, this concentration was used for subsequent analyses.

**Figure 4.**
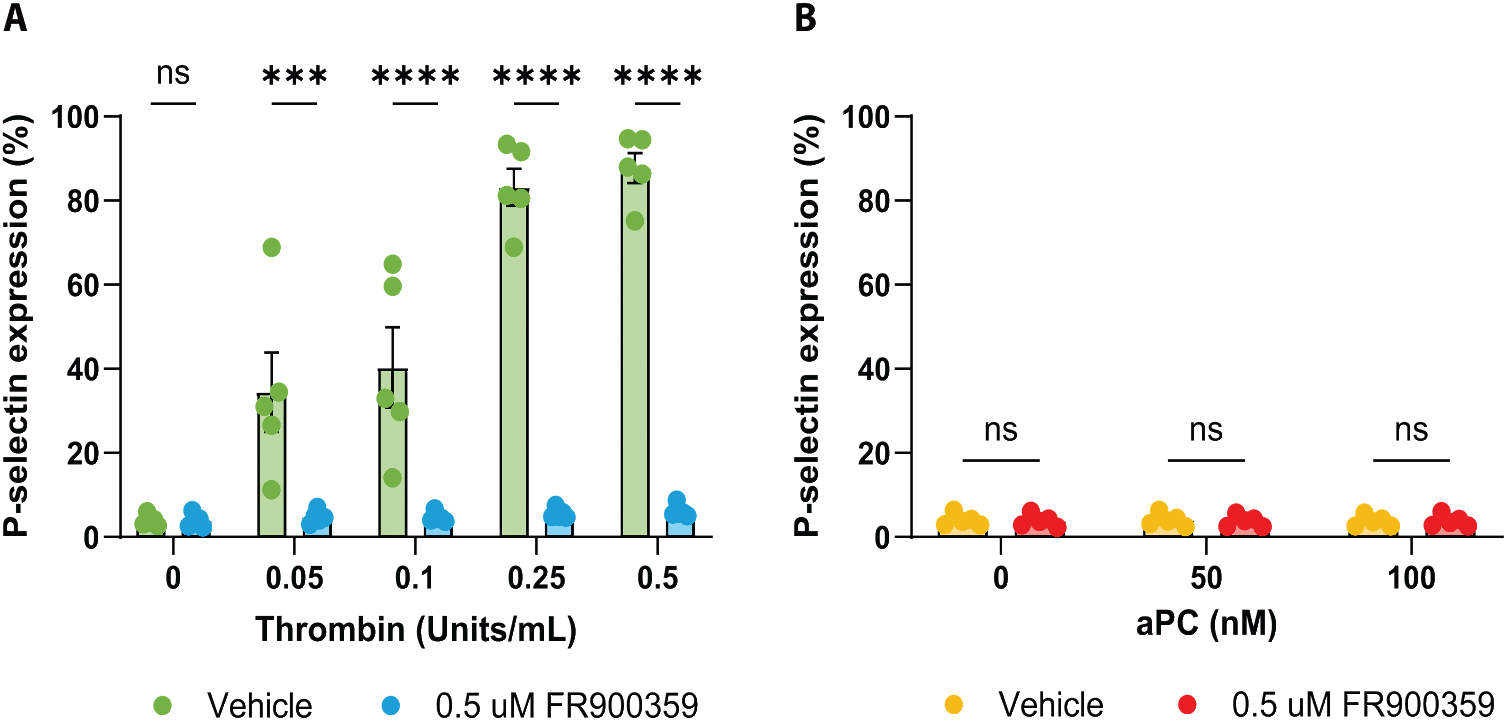
Protease-dependent Gαq signaling drives thrombin-induced but not aPC-induced platelet activation. **A)** Thrombin-induced P-selectin expression in human platelets in the presence of vehicle or 0.5 µM Gαq inhibitor. Thrombin induces a dose-dependent upregulation in P-selectin expression, which is significantly suppressed by FR900359 at higher thrombin concentrations. **B)** aPC-induced P-selectin expression in human platelets in the presence of vehicle or 0.5 µM Gαq inhibitor. aPC does not induce selectin expression across the tested concentrations, and FR900359 has no effect. Data are shown as mean ± SEM from five independent platelet donors (N = 5, n = 1). Statistical analysis was performed using two-way repeated-measures ANOVA with Geisser-Greenhouse correction followed by Šídák’s multiple-comparisons test. Significance thresholds were denoted as *** for P ≤ 0.001 and **** for P ≤ 0.0001.

Two-way repeated-measures ANOVA revealed a significant interaction between thrombin concentration and Gαq inhibitor treatment, indicating that Gαq inhibition altered the thrombin dose-response relationship (**Supplementary Table S8**). Post hoc analysis showed that treatment with 0.5 µM FR900359 had no effect on baseline or low thrombin-induced responses but significantly suppressed P-selectin expression at higher thrombin concentrations (0.25 and 0.5), demonstrating a strong dependence on Gαq activity for thrombin-induced platelet activation under these conditions (**Figure 4A; Supplementary Table S9)**. Increasing FR900359 concentrations did not further suppress thrombin-induced responses (**Supplementary Figure S5A**). Assessment of integrin αIIb activation showed a similar directional pattern but did not reach statistical significance across inhibitor concentrations **(Supplementary Figure S5B)**. These platelet findings are consistent with the thrombin- and FR900359-sensitive PAR1 signaling branch identified in recombinant cell assays. We do not interpret NFκB1 reporter activity in HEK293 cells as a direct mechanistic driver of platelet P-selectin exposure. NFκB1 reporter activity and platelet activation serve here as distinct downstream readouts in different cellular contexts.

In contrast, activated protein C (aPC) failed to induce measurable P-selectin expression across the concentration range tested (**Figure 4B, Supplementary Figure S5C**). Consistent with this, two-way repeated-measures ANOVA revealed no interaction between aPC concentration and Gαq inhibitor treatment and no main effects of aPC concentration or Gαq inhibition (**Supplementary Table S8**). Post hoc testing confirmed that 0.5 µM FR900359 did not affect aPC-treated platelets at any concentration (**Supplementary Table S9)**. Integrin αIIb activation likewise remained unchanged in response to aPC and was unaffected by FR900359 treatment (**Supplementary Figure S5D**). This lack of platelet activation occurred despite evidence in our HEK293 assays that aPC can activate PAR1 and engage Gα12-linked signaling, suggesting that a Gα12-biased PAR1 state is not sufficient to drive these platelet activation markers in this experimental context.

To more directly assess proximal platelet signaling, we next measured calcium flux in calcium-free buffer conditions. Thrombin produced robust and concentration-dependent calcium responses at both 0.1 and 0.5 U/mL, and these responses were significantly reduced by FR900359 (2.5 µM), consistent with engagement of a Gαq-sensitive platelet signaling branch (**Supplementary Figure S6; Supplementary Table S10**). Two-way repeated-measures ANOVA showed significant effects of condition and significant time vs condition interactions for both 0.1 U/mL thrombin (condition: P = 0.0127; time vs condition: P < 0.0001) and 0.5 U/mL thrombin (condition: P = 0.0009; time vs condition: P < 0.0001). In contrast, simultaneous addition of extracellular calcium with 50 nM aPC did not produce a detectable increase in calcium flux above the response observed with calcium alone (condition: P = 0.7365; time vs condition: P > 0.9999) (**Supplementary Figure S6; Supplementary Table S10**). These data extend the P-selectin and integrin findings by showing that thrombin, but not aPC, engages a proximal platelet calcium readout.

Together, these platelet studies support that thrombin engages a Gαq-sensitive platelet signaling branch reflected in calcium flux and activation marker expression. In contrast, aPC did not measurably activate platelets or elicit a detectable calcium response above baseline in the presence of calcium alone, underscoring the context dependence of PAR1 functional selectivity and downstream effector coupling.

## DISCUSSION

Across assays, thrombin drives a PAR1 signaling state that engages both Gαq and Gα12, whereas aPC produces detectable coupling to Gα12 without detectable Gαq. Neither yielded detectable β-arrestin-2 recruitment in PRESTO-Tango under our assay conditions. This divergence is echoed downstream by NFκB1 versus THRB transcriptional reporters and by platelet activation markers that require Gαq activity under the conditions tested. Thus, in our system, PAR1 functional selectivity is organized primarily along a Gαq-versus-Gα12 axis within a largely G protein-biased receptor. Thrombin and aPC define two naturally occurring points along this axis, providing an endogenous model for dissecting functional selectivity. Here, we connect these protease-defined PAR1 signaling states to transcriptional programs and platelet phenotypes using complementary biosensor assays and primary human platelets.

By directly measuring 13 individual Gα subunits with TRUPATH, we extend earlier work that linked PAR1 signaling—and thrombin-triggered outputs in particular— to Gαq- and Gα12/13-associated pathways largely through downstream calcium, RhoA, permeability, and migration readouts rather than subtype-resolved, transducer-wide profiling of individual G proteins (Hung et al., 1992; Offermanns et al., 1994; Gavard and Gutkind, 2008; McCoy et al., 2010). Our findings are also consistent with more recent biosensor and structural studies indicating that PAR1 can engage multiple G-protein families, while adding a direct side-by-side comparison of thrombin and aPC at the level of individual Gα subunits (Avet et al., 2022; Guo et al., 2024). Thrombin produced detectable coupling to Gαq and Gα12, while aPC produced detectable coupling only to Gα12. Because sensor-to-receptor DNA ratios are optimized independently for each TRUPATH sensor, these data are most robustly interpreted as a qualitative divergence in detectable coupling rather than as absolute comparisons of coupling efficiency between Gα subunits. Nevertheless, the loss of detectable Gαq engagement with aPC provides a transducer-level explanation for protease-dependent divergence in downstream outputs.

Downstream, analysis of the PAR1 ± thrombin TRE-MPRA dataset from Zahm et al. and our targeted dual-luciferase assays nominate and validate TRE reporters that separate these branches. The NFκB1 TRE was induced by thrombin and abolished by FR900359, consistent with dependence on Gαq-family signaling. This pattern aligns with prior work showing that thrombin-activated PAR1 can signal through Gαq, Gβγ, phospholipase C, calcium, and protein kinase C to promote NFκB activation and inflammatory gene programs in endothelial contexts (Rahman et al., 2002; Gambaryan et al., 2010). In contrast, THRB was induced by both thrombin and aPC and was not reduced by FR900359; if anything, maximal THRB activity trended upward with FR900359 pretreatment in our dataset (**Supplementary Table S5**), suggesting that inhibiting Gαq-family signaling can shift net transcriptional output toward FR900359-insensitive pathways. However, additional work will be needed to map the intermediates linking PAR1-Gα12 engagement to THRB activation in HEK293 cells.

Notably, Zahm and colleagues pointed out that the THRB motif is highly similar to a CArG box within the serum response element bound by serum response factor (SRF), suggesting that this TRE may function primarily as a readout of SRF/MRTF-dependent transcription rather than THRB-specific nuclear receptor biology (Zahm et al., 2024). SRF/MRTF activity is tightly coupled to RhoA-regulated actin dynamics. Because GPCR-mediated activation of RhoA commonly occurs downstream of Gα12/13 (and can involve Gαq/11 in a context-dependent manner), and because PAR signaling in platelets engages RhoA downstream of Gαq and Gα12/13 (Vogt et al., 2003; Jin et al., 2009; Yu and Brown, 2015), the FR900359-insensitive activation of THRB in HEK293 cells is compatible with a model in which THRB reports a Gαq-independent, Gα12-linked transcriptional branch (a Gα12/13-RhoA-SRF/MRTF-like pathway) under the experimental conditions tested. This interpretation is further supported by the observation that aPC selectively activates PAR1-Gα12 coupling in TRUPATH yet still robustly induces THRB. However, direct perturbation of the Gα12/13-RhoA-SRF/MRTF axis will be required to establish strict dependence.

Finally, primary platelet assays bridge these molecular signatures to a physiologic context. With PAR4 blocked to restrict thrombin responses to PAR1, thrombin induced robust P-selectin upregulation, which was significantly suppressed by FR900359. Integrin αIIbβ3 activation showed a similar directional trend but did not reach statistical significance, and FR900359 did not produce statistically significant suppression under these conditions. In contrast, aPC did not elicit measurable P-selectin or integrin activation across the concentrations tested, regardless of FR900359. Together, these data support that Gαq-family activity is required for these thrombin-driven platelet activation markers under the conditions tested, whereas aPC-driven, Gα12-biased signaling is insufficient to engage these outputs in this assay.

Our mechanistic data fit within a broader literature on protease-activated receptor signaling by coagulation proteases in endothelial cells. Endothelial cells co-express multiple PARs and co-receptors such as thrombomodulin and endothelial protein C receptor, which integrate signals from thrombin, aPC, and other coagulation factors (Rezaie, 2014). In that context, cleavage of PAR1 at Arg41 by thrombin is generally associated with barrier-disruptive and pro-inflammatory signaling. In contrast, cleavage at Arg46 by aPC, often in an endothelial protein C receptor and thrombomodulin-dependent manner, favors cytoprotective responses, including barrier stabilization and anti-inflammatory gene expression (Ludeman et al., 2005; Mosnier et al., 2012; Bouwens et al., 2013; De Ceunynck et al., 2018; Stojanovski and Di Cera, 2025). These cytoprotective effects extend to in vivo models, where aPC and aPC variants acting through PAR1 confer protection in ischemic and inflammatory injury settings (Xiang et al., 2025). Our HEK/HTLA system does not include overexpressed endothelial protein C receptor or thrombomodulin, and we did not assess whether recombinant PAR1 localized to the membrane microdomains implicated in endothelial aPC signaling. Because endothelial aPC-PAR1 cytoprotective signaling has been linked to EPCR/thrombomodulin-supported cleavage and to specific membrane compartmentalization of PAR1, the absence of detectable β-arrestin-2 recruitment in PRESTO-Tango likely reflects the lack of these endothelial context determinants rather than an absolute inability of aPC to engage arrestin-dependent PAR1 signaling. The inclusion of TRAP-6 strengthens this interpretation by showing that even a non-proteolytic PAR1 agonist fails to produce detectable PRESTO-Tango β-arrestin recruitment in the same recombinant system. Together, these observations suggest that the PAR1 arrestin-negative phenotype we observe is strongly context-dependent and should not be taken as inconsistent with the endothelial literature on aPC cytoprotection (Soh and Trejo, 2011). Nevertheless, the Gα12-associated transcriptional branch we define with THRB provides a candidate pathway that could contribute to aPC-driven cytoprotection in settings where Gαq-dependent inflammatory signaling is suppressed.

These findings also connect to recent structural and transducer-wide work on protease-activated receptors. Cryo-electron microscopy structures of PAR2 in complex with Gαq and Gα13 have shown how different tethered ligands can stabilize distinct active conformations and determine G protein selectivity at the structural level (Zhu et al., 2025). Together with EMTA coupling maps (Avet et al., 2022), these structures provide a transducer-wide and atomistic view of how GPCRs sample G protein space. Our functional PAR1 data suggest that distinct tethered ligands generated by thrombin and aPC drive the receptor into different regions of this space, favoring either a Gαq-plus-Gα12 state or a Gα12-selective state. Future structural work on PAR1 bound to thrombin- and aPC-generated tethered peptides, ideally in complex with Gαq and Gα12 heterotrimers, would test this idea directly and could reveal the conformational determinants of the Gαq versus Gα12 bias we observe.

Several limitations should be noted. Many of our assays rely on heterologous overexpression in HEK293 and HTLA cells, which may not capture native receptor density, PAR crosstalk, co-receptor expression, or protease localization. Furthermore, we did not assess membrane microdomain localization of recombinant receptors in HEK293/HTLA cells, so the present study does not address the contribution of lipid rafts to the observed signaling profiles in these assay systems. Our transcriptional analysis also relies on a TRE-MPRA dataset collected at a single stimulation duration and within a defined promoter architecture, and our targeted luciferase assays likewise report a single time point. In addition, PAR1 cleavage by aPC can be slower and less efficient than thrombin in endothelial settings (Ludeman et al., 2005), and co-factors such as EPCR and thrombomodulin can shape protease selectivity; therefore, additional coupling or kinetics could emerge outside our acquisition window or in other cellular contexts. The TRAP-6 control partially addresses this issue by showing that a non-proteolytic PAR1 agonist also produces detectable Gαq and Gα12 coupling without detectable PRESTO-Tango β-arrestin recruitment under the same assay conditions, supporting a contribution of receptor-state selectivity in addition to protease-specific activation kinetics. We also did not quantify endogenous EPCR or thrombomodulin expression in HEK293 or HTLA cells, nor did we directly determine whether aPC cleaved PAR1 at Arg41, Arg46, or another site under our assay conditions. Accordingly, our aPC data should be interpreted as defining the signaling profile generated in these recombinant systems rather than as direct evidence of Arg46 cleavage or of the full endothelial cytoprotective signaling program. Finally, the absence of detectable β-arrestin-2 recruitment by PRESTO-Tango should be interpreted cautiously, as PAR1’s proteolytic activation mechanism and receptor engineering may limit sensitivity. Future work incorporating endogenously relevant co-receptors and orthogonal arrestin readouts will be important to generalize these conclusions beyond the present assay conditions.

Despite these caveats, our data support PAR1 as a tractable model of endogenous functional selectivity that operates along a Gαq versus Gα12 axis within a largely G protein-biased receptor. Thrombin and aPC define two naturally occurring signaling states in this context: a Gαq-plus-Gα12 state associated with NFκB1 activation and thrombin-induced platelet P-selectin upregulation, and a Gα12-biased state that preserves THRB reporter output while failing to engage these platelet activation markers. The framework we apply here—linking proximal coupling profiles to transcriptional reporters and primary-cell phenotypes—can be extended to other GPCRs to help map and exploit functional selectivity.

## MATERIALS AND METHODS

### Reagents

DMEM was purchased from Thermo Fisher Scientific (Cat# 10566024). FBS (Cat# FB-11) and dialyzed FBS (Cat# FB-03) were purchased from Omega Scientific. Dopamine was purchased from Tocris Bioscience (Cat# 3548). Neurotensin (8-13) (trifluoroacetate salt) (Cat# 24718) and FR900359 (Cat# 33666) were purchased from Cayman Chemical. Thrombin was purchased from MedChemExpress (Cat# HY-114164). Activated Protein C (aPC) was purchased from Enzyme Research Laboratories (Cat# APC). TRAP-6 was purchased from TargetMol (Cat# T7625). All other chemicals were purchased from Sigma-Aldrich. Dilutions of stock drug solutions were made in 3X drug assay buffer (0.3 mg/mL ascorbic acid, 0.3% bovine serum albumin, and 20 mM HEPES in HBSS) or 6X for FR900359 assays. For PRESTO-Tango, dilutions of drug stocks were made in 5X drug assay buffer.

### Pharmacological Inhibitors

FR900359 (Cayman, Cat# 33666) is a macrocyclic depsipeptide originally isolated from Ardisia crenata that acts as a highly selective inhibitor of the Gαq family of G proteins (Kukkonen, 2016; Hermes et al., 2021; Voss, 2023). It binds to Gαq/11/14 and stabilizes the GDP-bound inactive state, thereby preventing receptor-stimulated GDP/GTP exchange and downstream signaling.

### Cell Culture

HEK293 (Cat# CRL-1573, RRID:CVCL_0045) cells were acquired from American Type Culture Collection (ATCC). HEK293 cells were maintained in DMEM (4.5 g/L D-glucose) supplemented with 10% FBS, penicillin (100 IU/mL), and streptomycin (100 µg/mL) (hereafter HEK growth media). HTLA cells were a gift from the Tao Che Lab and were maintained in DMEM supplemented with 10% FBS, 2 μg/mL puromycin, and 100 μg/mL hygromycin B (hereafter HTLA growth media). For relevant experiments, cells were transitioned to DMEM (4.5 g/L D-glucose) supplemented with 1% dialyzed FBS, penicillin (100 IU/mL), and streptomycin (100 µg/mL) (hereafter dFBS media).

### TRUPATH Assay

The TRUPATH biosensor platform (Olsen et al., 2020) is a resonance energy transfer-based system that enables direct measurement of GPCR coupling to individual Gα subunits. HEK293 cells were seeded at a density of 2 x 10^6^ cells per 10-cm plate in HEK growth media and incubated overnight (<24 hours) at 37°C in a humidified atmosphere containing 5% CO_2_ and >90% humidity. The TransIT-2020 reagent (Mirus Bio, Cat# MIR 5404) was equilibrated to room temperature for 30 minutes before use and gently vortexed immediately before preparing the transfection complexes. DNA mixtures were prepared using sensor-specific sensor-to-receptor ratios (sensor:receptor DNA) that were empirically optimized for each TRUPATH sensor to maximize BRET dynamic range. For Gαi3, GαoA, and Gα12, a 1:1 ratio was employed with 1480 ng of sensor and 1480 ng of receptor. For Gαz, a 2:1 ratio was used with 1480 ng of sensor and 740 ng of receptor. For all other sensors, a 5:1 ratio was applied with 1480 ng of sensor and 296 ng of receptor. Receptor plasmids encoding PAR1 or reference GPCRs (NTSR1, OPRK, and ADRB2) were utilized. Opti-MEM (Gibco, Cat# 31985070) was added at a ratio of 1 μL per 10 ng of DNA, and TransIT-2020 was added at 3 μL per 1 μg of DNA according to the manufacturer’s instructions. The DNA/Opti-MEM/TransIT-2020 mixtures were gently mixed by flicking and incubated at room temperature for 20 minutes to allow complex formation. The mixtures were then added dropwise to the 10-cm plates and evenly distributed by gently rocking the plates side to side. Transfected cultures were incubated overnight at 37°C, 5% CO_2_, and >90% humidity.

Prior to harvesting, cells were examined under an epifluorescence microscope to assess morphology, density, and transfection efficiency via GFP expression. Media was aspirated, and the cells were washed with 1 mL of Versene (Dulbecco’s phosphate-buffered saline containing 500 μM EDTA). After aspirating the wash, 3 mL of Versene was added, and the plate incubated for 3 minutes to facilitate detachment. Cells were gently dislodged by pipetting, collected by centrifugation at 300 x g for 3 minutes, and resuspended in 1 mL of dFBS media. The cell suspension was diluted to 10,000 cells per 40 μL (20,000 cells per 40 μL for Gs-short) as determined using a Countess™ 3 FL Automated Cell Counter (ThermoFisher, Cat# AMQAF2000, RRID:SCR_026963). Cells were plated in poly-D-Lysine-coated, clear-bottom 384-well assay plates (Greiner Bio-One, Cat# 781098; Sigma, Cat# P6407-5MG) at a density of 10,000 cells per well (20,000 cells per well for Gs-short). Plates were briefly centrifuged at 100 x g for 15-30 seconds to settle the cells and remove droplets from the sidewalls, then incubated for 24 hours at 37°C, 5% CO_2_, and >90% humidity.

Assay buffer (20 mM HEPES in 1x HBSS, pH 7.4 with KOH) and 3x drug buffer were prepared at least 2 hours prior to experimentation. Drugs were diluted to 3x working concentration in 3x drug buffer using 96-well plates (Fisher, Cat# 12-566-120) and serially diluted in log or half-log concentrations as appropriate (e.g., 10 μM, 3 μM, 1 μM, 0.3 μM) based on previously published potencies and adjusted as necessary for each experimental condition. The bioluminescent substrate Prolume Purple (NanoLight, Cat# 369) was prepared in Nanofuel (NanoLight, Cat# 399) to 1 mM and stored at −80°C. For the assay, growth media was removed from each well, and a white backing (PerkinElmer, Cat# 6005199) was applied to the plate bottom. Prolume Purple was diluted in assay buffer to 7.5 μM, and 20 μL was added to each well, followed by 10 μL of the drug solution.

Plates were read using a PHERAstar *FSX* (BMG Labtech, RRID:SCR_027001) with multichromatic, simultaneous dual-emission settings. Well scanning was performed using a 3 mm spiral average scan with the top optic. General instrument settings were a 0.1-second settling time, four kinetic cycles, and a 1-second pause between cycles. For each kinetic cycle, each individual well was sampled continuously for 0.5 seconds (the measurement interval for the spiral read) with the first read of each cycle beginning at 0 seconds (no initial delay). A full-plate kinetic cycle read required approximately 335 seconds. Gain settings were 3800 for Gain A and 3000 for Gain B, with a 40% gain adjustment applied to both channels. All wells were read regardless of occupancy. For concentration-response curve fitting, BRET ratios from the third kinetic cycle were used unless otherwise indicated.

### PRESTO-Tango Assay

The PRESTO-Tango system (Kroeze et al., 2015) was utilized to screen PAR1 ligand-receptor activation via the G protein-independent β-arrestin recruitment pathway. HTLA cells were plated on 10 cm tissue culture-treated dishes at a density of 3×10^6^ cells in HTLA growth media, then incubated for 6 hours at 37°C, 5% CO_2_, and >90% humidity. After 6 hours, cells were transfected with 5 μg of F2R-Tango plasmid (Addgene, Cat# 66276, RRID:Addgene_66276) or the reference DRD2-Tango plasmid (Addgene, Cat# 66269, RRID:Addgene_66269) using TransIT-2020 according to the manufacturer’s instructions.

The next day, cells were detached from culture dishes with Versene (Dulbecco’s phosphate-buffered saline containing 500 μM EDTA), collected by centrifugation at 300 x g for 3 minutes, and resuspended in 1 mL of dFBS media. The concentration of the cell suspension was determined using a Countess™ 3 FL Automated Cell Counter (ThermoFisher, Cat# AMQAF2000, RRID:SCR_026963) and then diluted to 15,000 cells per 40 μL. Cells were plated in poly-D-Lysine-coated, clear-bottom 384-well assay plates (Greiner Bio-One, Cat# 781098; Sigma, Cat# P6407-5MG) at a density of 15,000 cells per well. Plates were briefly centrifuged at 100 x g for 15-30 seconds, then incubated for 24 hours at 37°C, 5% CO_2_, and >90% humidity.

The following day, 10 µL of 5X drug dilutions in drug assay buffer were added to wells, and plates were incubated overnight. Luciferase activity was measured on a PHERAstar *FSX* (BMG Labtech, RRID:SCR_027001) using the Bright-Glo Luciferase Assay (Promega, Cat# E2610). Bright-Glo reagent was diluted in an equal volume of assay buffer (20 mM HEPES in HBSS), and 10 μL of diluted reagent was added to each well of the plate.

### TGFα Shedding Assay

TGFα shedding assay was performed as described previously (Inoue et al., 2012, 2019). HEK293 cells were plated in a 6-well tissue culture-treated plate at 5×10^5^ cells per well and incubated overnight at 37°C, 5% CO_2_, and >90% humidity. The next day, cells were transfected with 500 ng of AP-TGFα-encoding plasmid and 200 ng of PAR1 or NTSR1 receptor plasmid using TransIT-2020 according to the manufacturer’s instructions and incubated overnight.

The following day, cells were detached from 6-well plates with Versene (Dulbecco’s phosphate-buffered saline containing 500 μM EDTA), washed in HBSS containing 5 mM HEPES (pH 7.4) (hereafter TGFα assay buffer), and plated on 96-well tissue culture-treated plates (30,000 cells/well in 90 µL of TGFα assay buffer). 30 minutes after seeding in the 96-well plate, 10 µL of 10X drug dilutions in TGFα assay buffer with 0.01% bovine serum albumin were added to the cells. The plate was incubated for 1 hour. After ligand incubation, plates were centrifuged at 190 x g for 2 min. Conditioned media (80 µL/well) was transferred to a fresh 96-well plate and equilibrated to room temperature. Both the conditioned media plate and the original cell plate (containing cells and residual media) were supplemented with 80 µL/well of p-NPP substrate (10 mM) (GoldBio, Cat# N-380-50) in AP buffer (120 mM Tris-HCl (pH 9.5), 40 mM NaCl, 10 mM MgCl_2_ in DI water). Plates were incubated at room temperature for 5 minutes before an initial absorbance measurement at 405 nm (OD405). After a further 1-hour incubation at room temperature, the absorbance at 405 nm was remeasured.

For the TGFα shedding assay FR900359 Gαq inhibition studies, 5 μL of a 20X FR900359 dilution was added to the corresponding wells and the plate incubated for 20 minutes. Subsequently, 5 μL of 20X drug dilution was added to the wells.

### TRE-MPRA Analysis

Publicly available TRE-MPRA data from Zahm et al. (2024) were obtained from the NCBI Gene Expression Omnibus (series accession GSE271608) and the accompanying Source Data tables. We extracted the PAR1 (F2R) dataset comparing thrombin stimulation versus vehicle (TRE-MPRA was performed only for PAR1 ± thrombin; aPC was assessed in subsequent targeted assays). Within the highest-responding promoter architecture (set 1, rotation 8, minCMV), TREs were ranked by the magnitude of thrombin-induced change (absolute log2 fold change) and statistical support, and the top 10 responses (largest positive and negative changes) were highlighted as candidate thrombin-responsive elements for follow-up. NFκB1 and THRB were selected from this set for targeted luciferase validation and mechanistic interrogation with FR900359.

### Dual-Glo Luciferase Assay

HEK293 cells were plated on 10 cm tissue culture-treated dishes at a density of 2×10^6^ cells in HEK growth media. The following day, cells were transfected with 1 µg of TRE reporter plasmid (TRE-driven Luc2P firefly luciferase with an internal SV40-driven Renilla luciferase control) and 1 µg of PAR1 receptor plasmid using TransIT-2020 according to the manufacturer’s instructions. The next day, cells were detached from culture dishes with Versene (Dulbecco’s phosphate-buffered saline containing 500 μM EDTA), washed in dFBS media, and plated on poly-D-Lysine-coated, clear-bottom 384-well assay plates (Greiner Bio-One, Cat# 781098; Sigma, Cat# P6407-5MG) at 20,000 cells/well in 20 µL of dFBS media.

The following day, 10 µL of drug dilutions in drug assay buffer were added to the wells, and the plates were incubated for six hours. Firefly and *Renilla* luciferase activities were then measured sequentially on a PHERAstar *FSX* (BMG Labtech, RRID:SCR_027001) using the Dual-Glo Luciferase Assay (Promega, Cat# E2920) according to the manufacturer’s instructions. Firefly luminescence was normalized to Renilla luminescence for each well.

For FR900359 Gαq inhibition studies, 5 μL of the 6X FR900359 dilution was added to the corresponding wells and the plate incubated for 20 minutes. Subsequently, 5 μL of 6X drug dilution was added to the wells.

### P-selectin and Integrin Expression in Platelets

Whole blood was collected under IRB 202408024, which was approved by the Washington University Institutional Review Board. The whole blood was centrifuged at 100 x g for 20 minutes at room temperature to generate platelet-rich plasma (PRP). PRP was supplemented with PGE1, then centrifuged at 500 x g for 20 minutes. The platelet pellet was washed by resuspending the pellet in Pipes Saline Glucose (PSG; 5 mM PIPES, 145 mM NaCl, 4 mM KCl, 50 μM Na_2_HPO_4_, 1 mM MgCl_2_, and 5.5 mM Glucose, pH 6.8) and supplemented with PGE1. The platelets were then centrifuged at 500 x g for 20 minutes. After the final centrifugation, the platelet pellet was resuspended in PSG before being diluted in M199 for activation assays. Washed platelets were incubated with BMS-986120 (400 nM, final) for 15 minutes at 37°C to inhibit PAR4, and ensure platelet activation was exclusively through PAR1 (Andrianova et al., 2026). Washed platelets (1×10^6^ total) in M199 media were stained with anti-human CD41a allophycocyanin, anti-human P-selectin PE, and PAC-1 FITC in the presence of thrombin (0.05, 0.1, 0.25, or 0.5 U/mL, final) or activated protein C (50 nM or 100 nM, final) for 15 minutes at 37°C. Platelet samples were fixed with BD FACS™ Lysing solution (BD, USA) and measured using CytoFlex (Beckman Coulter, USA, RRID:SCR_019627). Platelet activation studies were performed using samples from five independent human donors, with each donor contributing one biological replicate per condition.

### Platelet Calcium Flux Assay

Washed platelets were prepared as described above and resuspended in PSG at 2 x 10^8/mL. Platelets were loaded with 5 µM Fura-2 AM (ThermoFisher, Cat# F1221) for 30 minutes at 37°C. After 30 minutes, platelets were centrifuged at 500 x g for 20 minutes in the presence of PGE1. Platelets were then resuspended in Tyrode’s Buffer (0.137 M NaCl, 2.68 mM KCl, 0.485 mM NaH2PO4, 12 mM NaHCO3, 5 mM HEPES, 5.5 mM Glucose and 0.35% BSA; pH 7.35) without calcium or magnesium at 4 x 10^8^/mL. Platelets were then incubated with BMS-986120 (400 nM, final) to inhibit PAR4 in the presence or absence of FR900359 (2.5 µM, final) for 15 minutes. Baseline calcium levels were measured at 37°C using a FlexStation 3 (Molecular Devices, San Jose, CA) at 340 nm and 380 nm. After stabilization of the baseline readings for 45 seconds, the indicated agonists and calcium (2 mM, final) were added, and cytoplasmic calcium flux was measured for 10 minutes for all agonists. Calcium responses were quantified as the Fura-2 ratio (340/380) from four independent human donors, with each donor contributing one biological replicate per condition.

### Statistics

All analyses were performed in GraphPad Prism 10.6.1 (GraphPad, RRID:SCR_002798). Heatmaps were generated using GraphPad Prism. Dose-response curves were fit using three- or four-parameter nonlinear regression models. Response amplitude is reported as the span (Top - Bottom) parameter from the fitted model, which reflects the net ligand-induced change in signal independent of baseline. Symbols and error bars represent mean values and standard error of the mean (SEM) of the indicated numbers of independent experiments, respectively. No statistical method was used to predetermine sample size.

Platelet activation data (P-selectin and integrin αIIb activation) from independent donors (n = 5) were analyzed using a two-way repeated-measures ANOVA, with agonist concentration and FR900359 Gαq inhibitor treatment as within-subject factors, and donor identity as a repeated-measures factor (matched by subject). Sphericity was not assumed, and Geisser-Greenhouse correction was applied where appropriate. When significant interactions were detected, post hoc multiple comparisons were performed using Šídák’s multiple comparisons test. All inhibitor concentrations were included in the statistical models. For clarity, only the lowest inhibitor concentration that produced maximal inhibition (0.5 µM) is shown in the main figure, while data for higher concentrations are provided in the Supplementary Information.

Calcium flux time-course data from independent donors (n = 4) were analyzed using two-way repeated-measures ANOVA with time and treatment condition as within-subject factors, matched by both donor (stacked) and time (repeated measures across rows). Separate analyses were performed for 0.1 and 0.5 U/mL thrombin conditions (± FR900359) and for calcium versus calcium with aPC conditions. When significant main effects or time vs condition interactions were detected, post hoc comparisons were performed using Šídák’s multiple comparisons test at individual time points. Statistical significance was defined as P < 0.05.

## DATA AVAILABILITY STATEMENT

The original contributions presented in the study are included in the article/Supplementary Material; further inquiries can be directed to the corresponding authors.

## AUTHOR CONTRIBUTIONS

B.S.F. designed the study, performed experiments, analyzed data, and wrote the manuscript. R.A.C. performed P-selectin, integrin and calcium experiments in platelets, analyzed platelet data, and edited the manuscript. J.G.E. designed the study, analyzed data, and edited the manuscript.

## ACKNOWLEDGMENTS

We thank the Tao Che laboratory (Washington University in St. Louis) for generously providing HTLA cells used in PRESTO-Tango assays. We also thank Ayumi Inoue (Tohoku University) for his gift of the TGFα shedding assay plasmid and associated protocol.

