## Supplementary Material for "Protease-Activated Receptor 1 as an Endogenous Model of Peptidergic Gαq-Gα12-Biased G Protein Signaling"

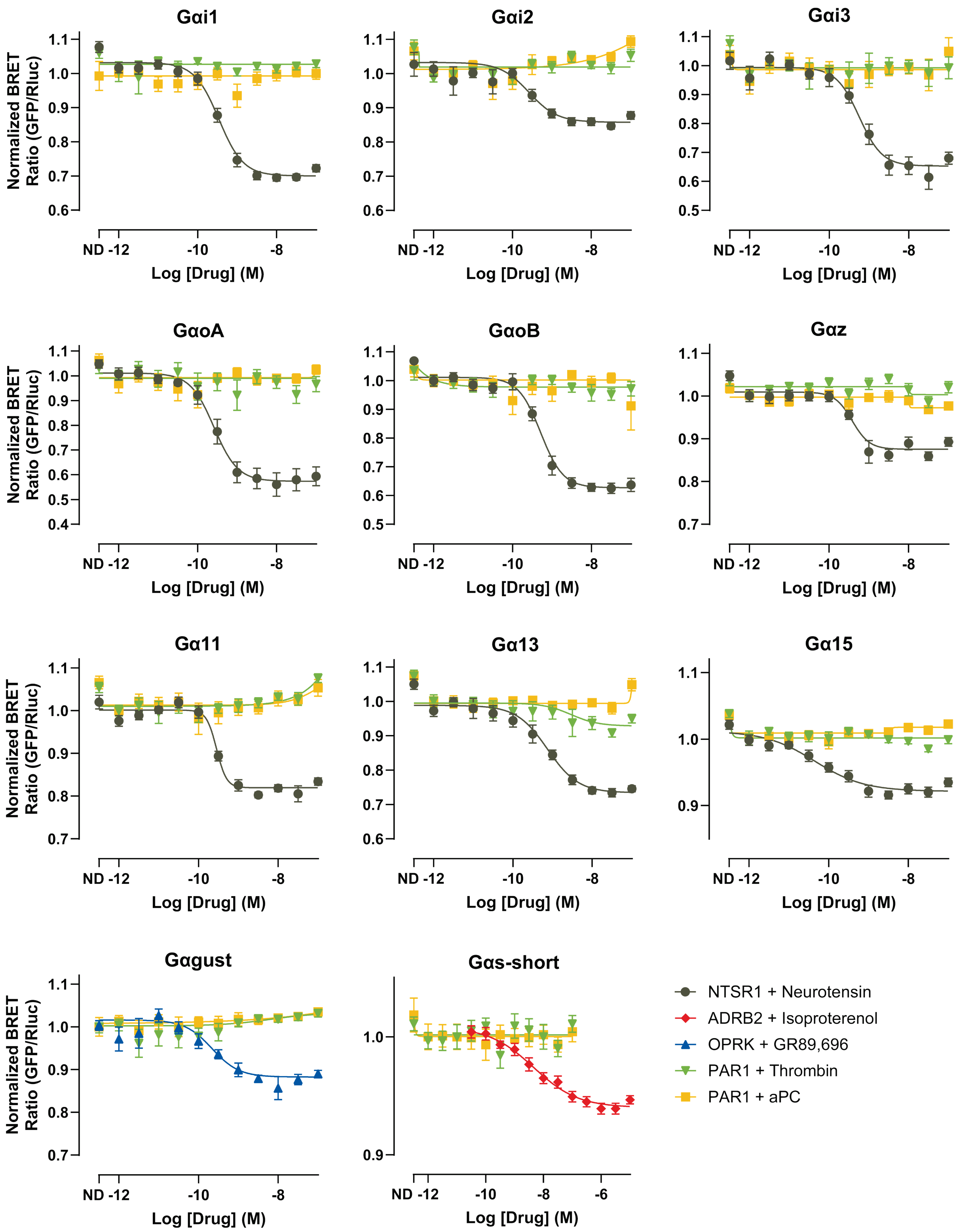


**Supplementary Figure S1. TRUPATH profiling reveals no detectable PAR1 coupling to non-Gαq/Gα12 subunits in response to thrombin or activated protein C (aPC).** TRUPATH concentration-response curves showing PAR1-mediated activation of the indicated Gα subunits following stimulation with thrombin or activated protein C (aPC). For each Gα subunit, a reference control receptor with established coupling was included to benchmark assay performance. Data are presented as mean ± SEM from 3 independent experiments with 4 technical replicates. Curves were fit using a four-parameter logistic model.

**Supplementary Table S1. TRUPATH Analysis of G Protein Coupling Profiles for NTSR1, PAR1, ADRB2, and OPRK.**

| Receptor | Transducer | Ligand | N,n | Log(EC50) | EC50 | Span |
|---|---|---|---|---|---|---|
| **NTSR1** | Gαi1 | Neurotensin | 3,4 | -9.462 ± 0.053 | 0.345 nM | 0.331 ± 0.012 |
| **PAR1** | Gαi1 | Thrombin | 3,4 | NR | NR | NR |
| **PAR1** | Gαi1 | aPC | 3,4 | NR | NR | NR |
| **NTSR1** | Gαi2 | Neurotensin | 3,4 | -9.581 ± 0.112 | 0.262 nM | 0.175 ± 0.013 |
| **PAR1** | Gαi2 | Thrombin | 3,4 | NR | NR | NR |
| **PAR1** | Gαi2 | aPC | 3,4 | NR | NR | NR |
| **NTSR1** | Gαi3 | Neurotensin | 3,4 | -9.248 ± 0.094 | 0.564 nM | 0.340 ± 0.022 |
| **PAR1** | Gαi3 | Thrombin | 3,4 | NR | NR | NR |
| **PAR1** | Gαi3 | aPC | 3,4 | NR | NR | NR |
| **NTSR1** | GαoA | Neurotensin | 3,4 | -9.585 ± 0.090 | 0.259 nM | 0.473 ± 0.027 |
| **PAR1** | GαoA | Thrombin | 3,4 | NR | NR | NR |
| **PAR1** | GαoA | aPC | 3,4 | NR | NR | NR |
| **NTSR1** | GαoB | Neurotensin | 3,4 | -9.289 ± 0.059 | 0.514 nM | 0.384 ± 0.015 |
| **PAR1** | GαoB | Thrombin | 3,4 | NR | NR | NR |
| **PAR1** | GαoB | aPC | 3,4 | NR | NR | NR |
| **NTSR1** | Gαz | Neurotensin | 3,4 | -9.419 ± 0.087 | 0.381 nM | 0.134 ± 0.009 |
| **PAR1** | Gαz | Thrombin | 3,4 | NR | NR | NR |
| **PAR1** | Gαz | aPC | 3,4 | NR | NR | NR |
| **NTSR1** | Gαq | Neurotensin | 3,4 | -9.443 ± 0.053 | 0.360 nM | 0.429 ± 0.018 |
| **PAR1** | Gαq | Thrombin | 3,4 | -8.236 ± 0.138 | 5.813 nM | 0.119 ± 0.014 |
| **PAR1** | Gαq | aPC | 3,4 | NR | NR | NR |
| **NTSR1** | Gα11 | Neurotensin | 3,4 | -9.547 ± 0.045 | 0.284 nM | 0.182 ± 0.008 |
| **PAR1** | Gα11 | Thrombin | 3,4 | NR | NR | NR |
| **PAR1** | Gα11 | aPC | 3,4 | NR | NR | NR |
| **NTSR1** | Gα12 | Neurotensin | 3,4 | -9.304 ± 0.108 | 0.496 nM | 0.190 ± 0.013 |
| **PAR1** | Gα12 | Thrombin | 3,4 | -9.288 ± 0.069 | 0.514 nM | 0.239 ± 0.010 |
| **PAR1** | Gα12 | aPC | 3,4 | -8.609 ± 0.106 | 2.461 nM | 0.126 ± 0.010 |
| **NTSR1** | Gα13 | Neurotensin | 3,4 | -9.128 ± 0.080 | 0.744 nM | 0.245 ± 0.014 |
| **PAR1** | Gα13 | Thrombin | 3,4 | NR | NR | NR |
| **PAR1** | Gα13 | aPC | 3,4 | NR | NR | NR |
| **NTSR1** | Gα15 | Neurotensin | 3,4 | -10.347 ± 0.197 | 0.045 nM | 0.091 ± 0.011 |
| **PAR1** | Gα15 | Thrombin | 3,4 | NR | NR | NR |
| **PAR1** | Gα15 | aPC | 3,4 | NR | NR | NR |
| **ADRB2** | Gαs,short | Isoproterenol | 3,4 | -8.346 ± 0.222 | 4.512 nM | 0.068 ± 0.009 |
| **PAR1** | Gαs,short | Thrombin | 3,4 | NR | NR | NR |
| **PAR1** | Gαs,short | aPC | 3,4 | NR | NR | NR |
| **OPRK** | Gαgust | GR89.696 | 3,4 | -9.663 ± 0.117 | 0.217 nM | 0.113 ± 0.010 |
| **PAR1** | Gαgust | Thrombin | 3,4 | NR | NR | NR |
| **PAR1** | Gαgust | aPC | 3,4 | NR | NR | NR |

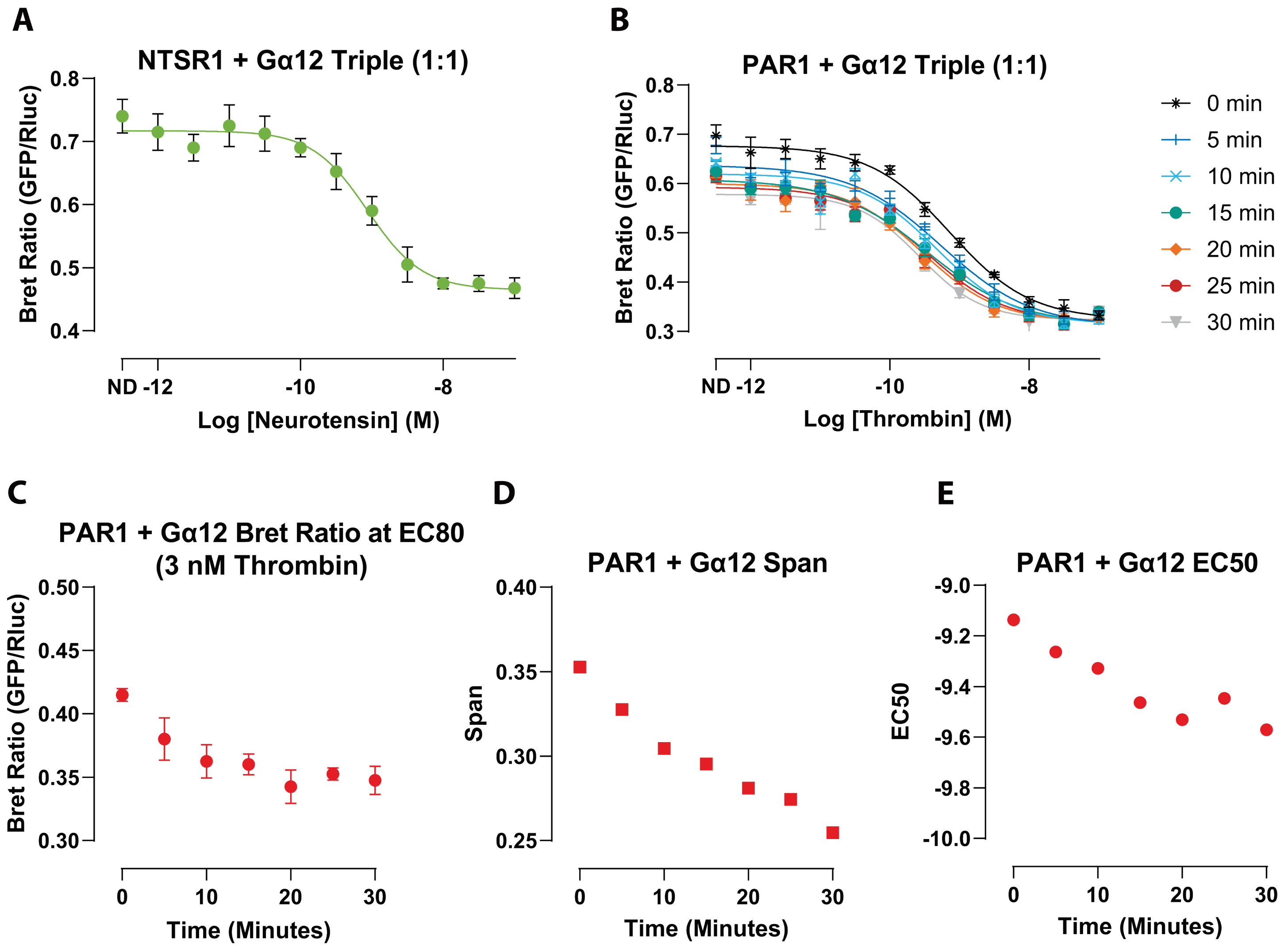


**Supplementary Figure S2. Time-Dependent TRUPATH Profiling of PAR1-Gα12 Signaling Following Thrombin Stimulation. A)** TRUPATH concentration-response curve for NTSR1-mediated activation of Gα12, shown as a reference control. **B)** TRUPATH concentration-response curves for PAR1-mediated activation of Gα12 following thrombin pre-incubation for increasing durations (0-30 min in 5-min intervals). **C-E)** Quantification of TRUPATH response parameters derived from the concentration-response curves shown in **(B)**, including BRET ratio **(C)**, signaling span **(D)**, and EC50 **(E)** as a function of thrombin incubation time. Data are presented as mean ± SEM from 3 independent experiments with 4 technical replicates. Concentration-response curves were fit using a four-parameter logistic model.

**Supplementary Table S2. TGFα Shedding Assay Quantification of NTSR1 and PAR1 Signaling with and without Gαq Inhibition.**

| Receptor | Ligand | Inhibitor | N,n | Log(EC50) | EC50 | Span |
|---|---|---|---|---|---|---|
| **NTSR1** | Neurotensin | None | 3,4 | -9.672 ± 0.062 | 0.21 nM | 28.935 ± 1.308 % |
| **PAR1** | Thrombin | None | 3,4 | -8.389 ± 0.099 | 4.08 nM | 16.389 ± 1.572 % |
| **PAR1** | Thrombin | FR900359 | 3,4 | -8.697 ± 0.105 | 2.01 nM | 10.314 ± 0.975 % |
| **PAR1** | aPC | None | 3,4 | -8.194 ± 0.073 | 6.39 nM | 20.016 ± 1.168 % |
| **PAR1** | aPC | FR900359 | 3,4 | -8.231 ± 0.191 | 5.88 nM | 25.176 ± 3.236 % |

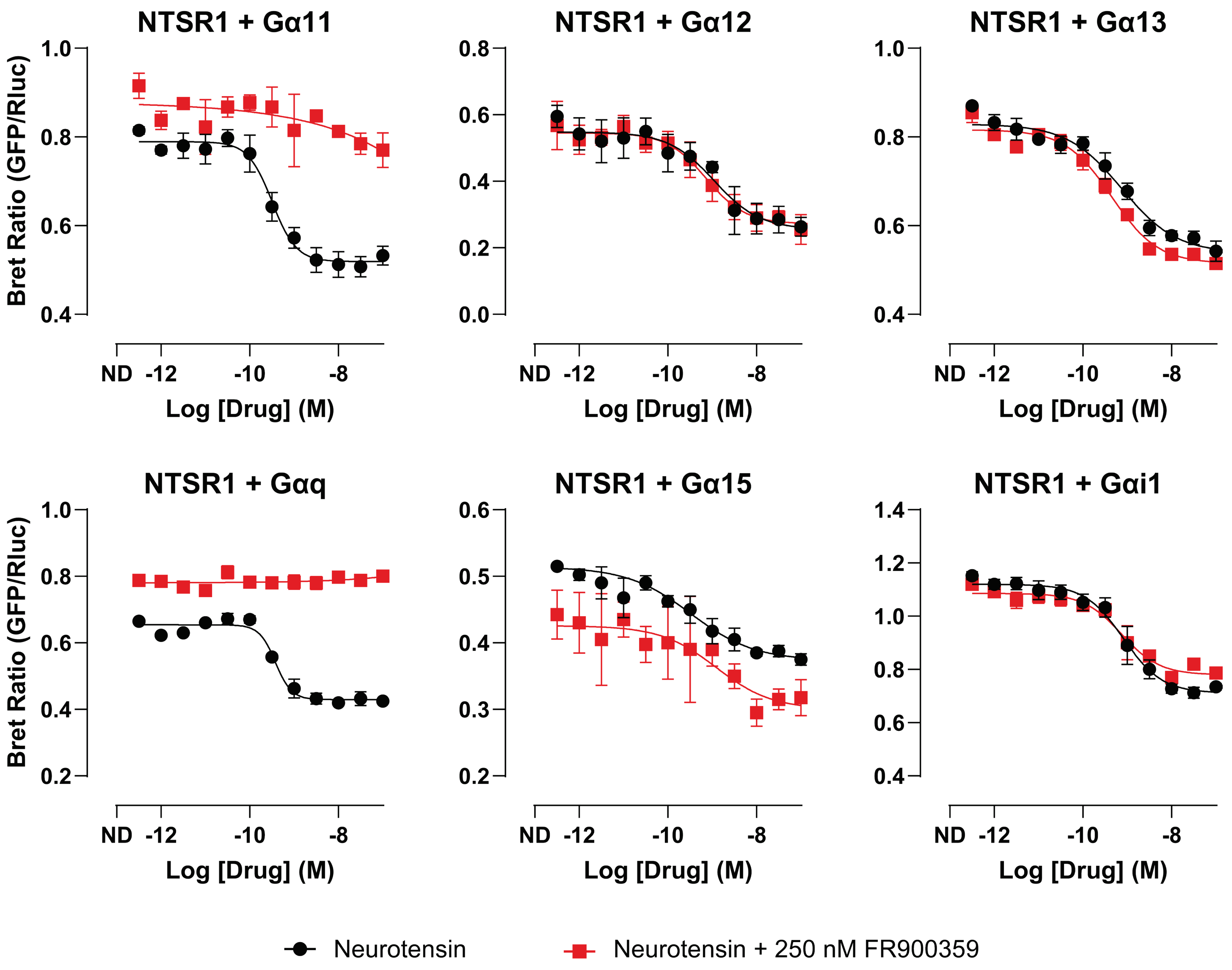


**Supplementary Figure S3. FR900359 abolishes Gαq/11 activation but preserves signaling through Gα12, Gα13, Gα15, and Gαi1 in the TRUPATH assay**. TRUPATH concentration–response curves for NTSR1-mediated activation of the indicated Gα subunits (Gα11, Gα12, Gα13, Gαq, Gα15, and Gαi1) in the presence or absence of the Gq inhibitor FR900359. FR900359 abolished detectable signaling through Gα11 and Gαq, while signaling via Gα12, Gα13, Gα15, and Gαi1 was preserved. Data are shown as mean ± SEM from 1 independent experiment with 4 technical replicates. Curves were fit using a four-parameter logistic model where responses were detectable.

**Supplementary Table S3. PRESTO-Tango β-Arrestin Recruitment Assay Fits for DRD2 and PAR1**

| Receptor | Ligand | N,n | Log(EC50) | EC50 | Span |
|---|---|---|---|---|---|
| **DRD2** | Dopamine | 3,4 | -8.439 ± 0.050 | 3.64 nM | 33.371 ± 0.763 |
| **PAR1** | Thrombin | 3,4 | NR | NR | NR |
| **PAR1** | aPC | 3,4 | NR | NR | NR |

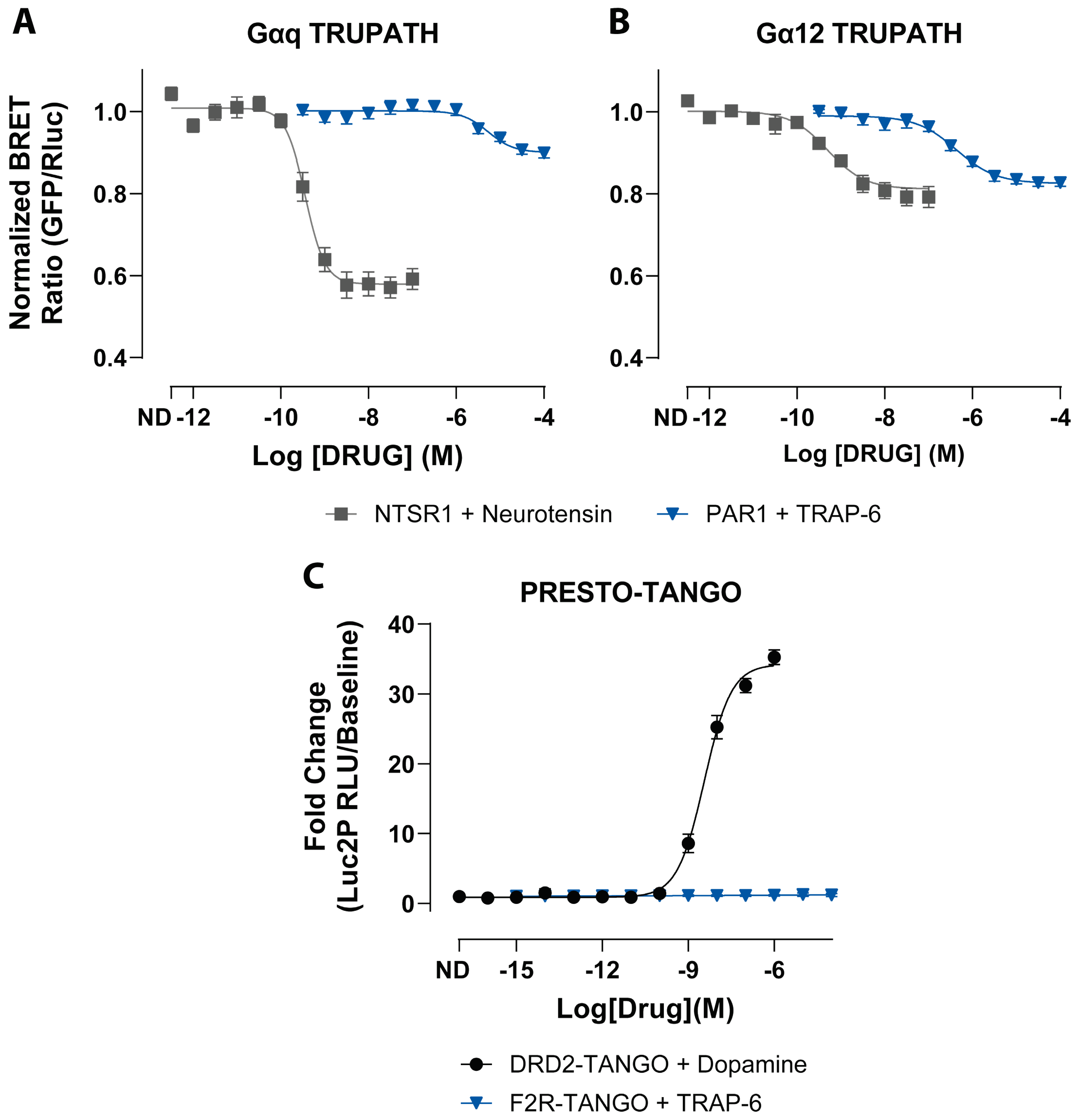


**Supplementary Figure S4. TRAP-6 activates PAR1-mediated G protein coupling but does not produce detectable β-arrestin recruitment in recombinant biosensor assays. A–B)** TRUPATH concentration-response curves for PAR1 stimulated with TRAP-6 (blue triangles) in Gαq **(A)** and Gα12 **(B)** biosensors. NTSR1 stimulated with neurotensin (gray squares) is shown as a reference control for assay performance. Under these assay conditions, TRAP-6 produced detectable PAR1 coupling to both Gαq and Gα12. **C**) PRESTO-Tango β-arrestin recruitment assay in HTLA cells. DRD2-Tango stimulated with dopamine (black circles) served as a reference control and produced robust, dose-dependent reporter activation, whereas F2R-Tango stimulated with TRAP-6 (blue triangles) did not produce a detectable increase in reporter signal across the tested concentration range. Data are presented as mean ± SEM from 3 independent experiments with 4 technical replicates. Concentration-response curves were fit using a four-parameter logistic model.

**Supplementary Table S4. Concentration-response fit parameters for TRAP-6 PAR1 signaling in TRUPATH and PRESTO-Tango assays**

| Assay | Receptor | Transducer | Ligand | N,n | Log(EC50) | EC50 | Span |
|---|---|---|---|---|---|---|---|
| **TRUPATH** | NTSR1 | Gα12 | Neurotensin | 3,4 | -9.304 ± 0.108 | 0.49 nM | 0.190 ± 0.013 |
| **TRUPATH** | NTSR1 | Gαq | Neurotensin | 3,4 | -9.443 ± 0.053 | 0.36 nM | 0.429 ± 0.018 |
| **TRUPATH** | PAR1 | Gα12 | TRAP-6 | 3,4 | -6.369 ± 0.110 | 0.43 µM | 0.165 ± 0.011 |
| **TRUPATH** | PAR1 | Gαq | TRAP-6 | 3,4 | -5.277 ± 0.161 | 5.29 µM | 0.102 ± 0.014 |
| **PRESTO-Tango** | DRD2 | β-arrestin-2 | Dopamine | 3,4 | -8.439 ± 0.050 | 3.64 nM | 33.371 ± 0.763 |
| **PRESTO-Tango** | PAR1 | β-arrestin-2 | TRAP-6 | 3,4 | NR | NR | NR |

**Supplementary Table S5. Transcriptional Reporter (TRE) Assay Fits for PAR1-Mediated NFκB1 and THRB Signaling**

| Receptor | Transducer | Ligand | Inhibitor | N,n | Log(EC50) | EC50 | Span |
|---|---|---|---|---|---|---|---|
| **PAR1** | NFkB1 | Thrombin | None | 3,4 | -8.319 ± 0.098 | 4.79 nM | 9.788 ± 0.706 |
| **PAR1** | NFkB1 | Thrombin | FR900359 | 3,4 | NR | NR | NR |
| **PAR1** | NFkB1 | aPC | None | 3,4 | NR | NR | NR |
| **PAR1** | NFkB1 | aPC | FR900359 | 3,4 | NR | NR | NR |
| **PAR1** | THRB | Thrombin | None | 3,4 | -8.023 ± 0.165 | 9.48 nM | 5.241 ± 0.536 |
| **PAR1** | THRB | Thrombin | FR900359 | 3,4 | -7.863 ± 0.281 | 13.72 nM | 8.939 ± 1.687 |
| **PAR1** | THRB | aPC | None | 3,4 | -7.856 ± 0.146 | 13.92 nM | 7.885 ± 0.776 |
| **PAR1** | THRB | aPC | FR900359 | 3,4 | -7.752 ± 0.152 | 17.72 nM | 9.512 ± 1.032 |

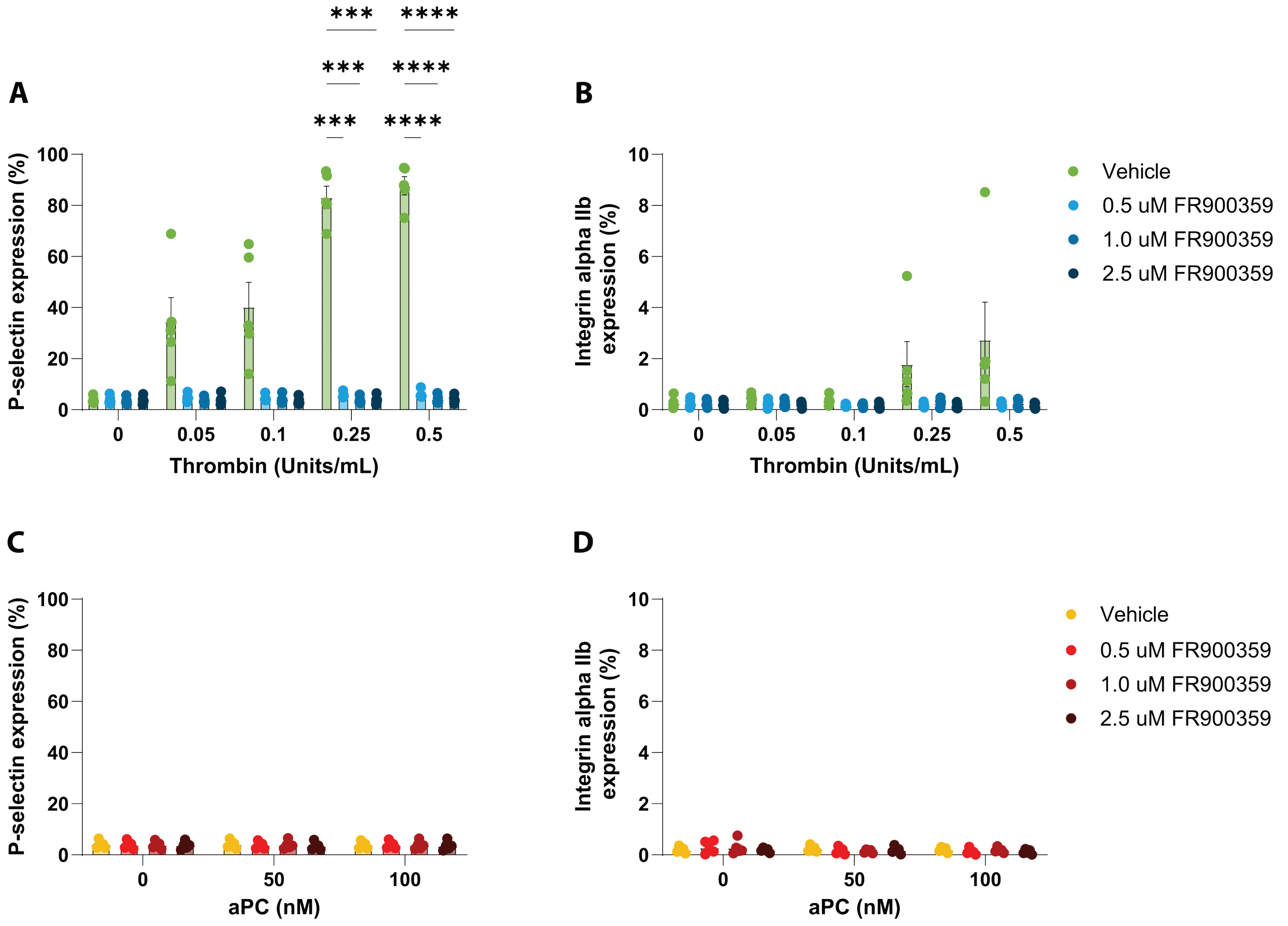


**Supplementary Figure S5. Higher concentrations of FR900359 Gαq inhibitor do not further suppress thrombin-induced platelet activation. A)** Thrombin-induced P-selectin expression in human platelets treated with vehicle or increasing concentrations of FR900359 (0.5, 1.0, and 2.5 µM). Increasing inhibitor concentration did not further suppress thrombin-induced P-selectin expression at any thrombin concentration, indicating near-maximal inhibition at the lowest concentration tested. **B)** Thrombin-induced integrin αIIb activation under the same conditions. No statistically significant differences were observed between vehicle- and FR900359-treated samples at any inhibitor concentration. **C)** aPC-induced P-selectin expression in the presence of vehicle or increasing concentrations of FR900359. aPC did not induce measurable P-selectin expression, and FR900359 treatment had no effect at any concentration tested. **D)** aPC-induced integrin αIIb activation in the presence of vehicle or FR900359. No differences were observed between vehicle and inhibitor-treated samples, consistent with the absence of aPC-induced platelet activation under these conditions. Data are shown as mean ± SEM from five independent platelet donors. Statistical analyses were performed using two-way repeated-measures ANOVA with Geisser-Greenhouse correction followed by Šídák's multiple-comparisons test. Significance thresholds were denoted as *** for $P \leq 0.001$ and **** for $P \leq 0.0001$.

**Supplementary Table S6. Šídák multiple-comparisons tests for all Gαq inhibitor concentrations for Thrombin-mediated P-selectin expression.**

| Thrombin Concentration (Units/mL) | Inhibitor Comparison | Mean Difference | 95% CI of Difference | Adjusted P Value | Summary |
|---|---|---|---|---|---|
| **0** | Vehicle vs 0.5 µM | 0.03 | -3.192 to 3.252 | >0.9999 | ns |
| **0** | Vehicle vs 1.0 µM | 0.222 | -2.665 to 3.109 | >0.9999 | ns |
| **0** | Vehicle vs 2.5 µM | 0.24 | -3.030 to 3.510 | >0.9999 | ns |
| **0** | 0.5 vs 1.0 µM | 0.192 | -3.051 to 3.435 | >0.9999 | ns |
| **0** | 0.5 vs 2.5 µM | 0.21 | -3.309 to 3.729 | >0.9999 | ns |
| **0** | 1.0 vs 2.5 µM | 0.018 | -3.271 to 3.307 | >0.9999 | ns |
| **0.05** | Vehicle vs 0.5 µM | 29.67 | -15.85 to 75.18 | 0.1925 | ns |
| **0.05** | Vehicle vs 1.0 µM | 30.65 | -14.93 to 76.24 | 0.1760 | ns |
| **0.05** | Vehicle vs 2.5 µM | 30.56 | -14.79 to 75.92 | 0.1763 | ns |
| **0.05** | 0.5 vs 1.0 µM | 0.986 | -1.994 to 3.966 | 0.8606 | ns |
| **0.05** | 0.5 vs 2.5 µM | 0.898 | -3.019 to 4.815 | 0.9695 | ns |
| **0.05** | 1.0 vs 2.5 µM | -0.088 | -3.910 to 3.734 | >0.9999 | ns |
| **0.1** | Vehicle vs 0.5 µM | 35.63 | -10.39 to 81.66 | 0.1158 | ns |
| **0.1** | Vehicle vs 1.0 µM | 36.22 | -9.662 to 82.10 | 0.1094 | ns |
| **0.1** | Vehicle vs 2.5 µM | 36.69 | -9.283 to 82.67 | 0.1055 | ns |
| **0.1** | 0.5 vs 1.0 µM | 0.588 | -2.774 to 3.950 | 0.9915 | ns |
| **0.1** | 0.5 vs 2.5 µM | 1.062 | -1.765 to 3.889 | 0.7868 | ns |
| **0.1** | 1.0 vs 2.5 µM | 0.474 | -2.990 to 3.938 | 0.9980 | ns |
| **0.25** | Vehicle vs 0.5 µM | 77.51 | 56.40 to 98.62 | 0.0003 | *** |
| **0.25** | Vehicle vs 1.0 µM | 79.17 | 58.12 to 100.2 | 0.0003 | *** |
| **0.25** | Vehicle vs 2.5 µM | 79.32 | 58.45 to 100.2 | 0.0002 | *** |
| **0.25** | 0.5 vs 1.0 µM | 1.66 | -1.089 to 4.409 | 0.3506 | ns |
| **0.25** | 0.5 vs 2.5 µM | 1.806 | -1.467 to 5.079 | 0.4227 | ns |
| **0.25** | 1.0 vs 2.5 µM | 0.146 | -3.194 to 3.486 | >0.9999 | ns |
| **0.5** | Vehicle vs 0.5 µM | 81.59 | 64.85 to 98.32 | <0.0001 | **** |
| **0.5** | Vehicle vs 1.0 µM | 83.62 | 66.91 to 100.3 | <0.0001 | **** |
| **0.5** | Vehicle vs 2.5 µM | 83.91 | 67.26 to 100.5 | <0.0001 | **** |
| **0.5** | 0.5 vs 1.0 µM | 2.03 | -1.329 to 5.389 | 0.3514 | ns |
| **0.5** | 0.5 vs 2.5 µM | 2.32 | -1.158 to 5.798 | 0.2623 | ns |
| **0.5** | 1.0 vs 2.5 µM | 0.29 | -3.235 to 3.815 | 0.9999 | ns |

**Supplementary Table S7. Šídák multiple-comparisons tests for all Gαq inhibitor concentrations for aPC-mediated P-selectin expression.**

| aPC Concentration (nM) | Inhibitor Comparison | Mean Difference | 95% CI of Difference | Adjusted P Value | Summary |
|---|---|---|---|---|---|
| **0** | Vehicle vs 0.5 µM | 0.172 | -2.994 to 3.338 | >0.9999 | ns |
| **0** | Vehicle vs 1.0 µM | 0.106 | -2.962 to 3.174 | >0.9999 | ns |
| **0** | Vehicle vs 2.5 µM | 0.32 | -2.876 to 3.516 | 0.9997 | ns |
| **0** | 0.5 vs 1.0 µM | -0.066 | -3.183 to 3.051 | >0.9999 | ns |
| **0** | 0.5 vs 2.5 µM | 0.148 | -3.091 to 3.387 | >0.9999 | ns |
| **0** | 1.0 vs 2.5 µM | 0.214 | -2.936 to 3.364 | >0.9999 | ns |
| **50** | Vehicle vs 0.5 µM | 0.272 | -2.863 to 3.407 | 0.9999 | ns |
| **50** | Vehicle vs 1.0 µM | 0.146 | -3.204 to 3.496 | >0.9999 | ns |
| **50** | Vehicle vs 2.5 µM | 0.628 | -2.664 to 3.920 | 0.9888 | ns |
| **50** | 0.5 vs 1.0 µM | -0.126 | -3.355 to 3.103 | >0.9999 | ns |
| **50** | 0.5 vs 2.5 µM | 0.356 | -2.807 to 3.519 | 0.9994 | ns |
| **50** | 1.0 vs 2.5 µM | 0.482 | -2.890 to 3.854 | 0.9976 | ns |
| **100** | Vehicle vs 0.5 µM | -0.062 | -2.970 to 2.846 | >0.9999 | ns |
| **100** | Vehicle vs 1.0 µM | -0.03 | -3.213 to 3.153 | >0.9999 | ns |
| **100** | Vehicle vs 2.5 µM | 0.202 | -3.247 to 3.651 | >0.9999 | ns |
| **100** | 0.5 vs 1.0 µM | 0.032 | -3.221 to 3.285 | >0.9999 | ns |
| **100** | 0.5 vs 2.5 µM | 0.264 | -3.235 to 3.763 | >0.9999 | ns |
| **100** | 1.0 vs 2.5 µM | 0.232 | -3.423 to 3.887 | >0.9999 | ns |

**Supplementary Table S8. Summary of two-way repeated-measures ANOVA results for platelet P-selectin expression.**

| Agonist | Factor | Effect Tested | P value | Interpretation |
|---|---|---|---|---|
| **Thrombin** | Thrombin concentration vs Gαq inhibitor | Interaction | < 0.0001 | Dose-response differs with Gαq inhibition |
| **Thrombin** | Thrombin concentration | Main effect | < 0.0001 | Robust dose-dependent activation |
| **Thrombin** | Gαq inhibitor treatment | Main effect | < 0.0001 | Strong suppressive effect |
| **Thrombin** | Subject (donor) | Random effect | < 0.0001 | Expected donor variability |
| **aPC** | aPC concentration vs Gαq inhibitor | Interaction | 0.532 | No interaction detected |
| **aPC** | aPC concentration | Main effect | 0.235 | No activation |
| **aPC** | Gαq inhibitor treatment | Main effect | 0.978 | No inhibitor effect |
| **aPC** | Subject (donor) | Random effect | < 0.0001 | Expected donor variability |

**Supplementary Table S9. Post hoc Šídák multiple-comparisons analysis of Gαq inhibitor effects on platelet P-selectin expression.**

| Agonist | Agonist Concentration | Comparison | Adjusted P value | Significance |
|---|---|---|---|---|
| **Thrombin** | 0 | Vehicle vs 0.5 μM Gαq inhibitor | > 0.9999 | Ns |
| **Thrombin** | 0.05 | Vehicle vs 0.5 μM Gαq inhibitor | 0.1925 | Ns |
| **Thrombin** | 0.1 | Vehicle vs 0.5 μM Gαq inhibitor | 0.1158 | Ns |
| **Thrombin** | 0.25 | Vehicle vs 0.5 μM Gαq inhibitor | 0.0003 | *** |
| **Thrombin** | 0.5 | Vehicle vs 0.5 μM Gαq inhibitor | < 0.0001 | **** |
| **aPC** | 0 nM | Vehicle vs 0.5 μM Gαq inhibitor | > 0.9999 | Ns |
| **aPC** | 50 nM | Vehicle vs 0.5 μM Gαq inhibitor | 0.9999 | Ns |
| **aPC** | 100 nM | Vehicle vs 0.5 μM Gαq inhibitor | > 0.9999 | Ns |

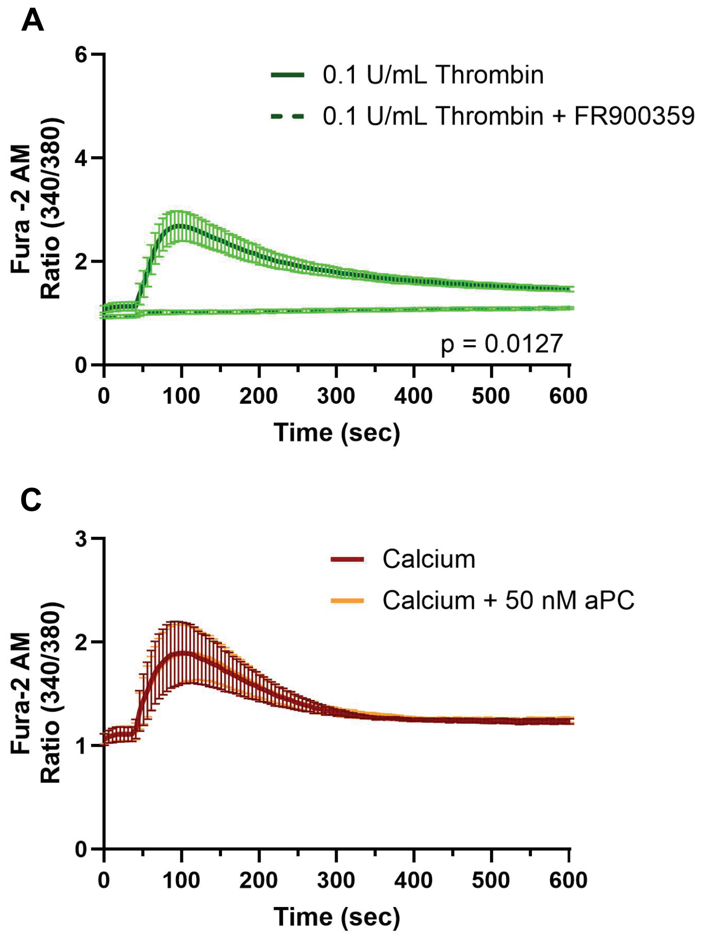


**Supplementary Figure S6. Platelet calcium flux reveals a thrombin-responsive, FR900359-sensitive signal, whereas aPC does not increase the response above calcium alone.** Washed human platelets were loaded with Fura-2 AM and assayed in calcium-free buffer. Platelets were incubated with BMS-986120 (400 nM, final) to restrict thrombin responses to PAR1. Where indicated, platelets were treated with FR900359 (2.5 µM, final). Calcium responses were recorded as the Fura-2 ratio (340/380 nm) over time from 4 independent human donors. **A)** Calcium flux induced by 0.1 U/mL thrombin in the presence or absence of FR900359. Condition and time vs condition effects were significant ($P = 0.0127$ and $P < 0.0001$, respectively). **B)** Calcium flux induced by 0.5 U/mL thrombin in the presence or absence of FR900359. Condition and time vs condition effects were significant ($P = 0.0009$ and $P < 0.0001$, respectively). **C)** Calcium-only and calcium + 50 nM aPC conditions, in which extracellular calcium and aPC were added simultaneously at the start of acquisition. No significant effect of condition was detected ($P = 0.7365$), and no significant time vs condition interaction was observed ($P > 0.9999$). Statistical analysis was performed using two-way repeated-measures ANOVA with Šídák's multiple comparisons test. Summary statistics are provided in Supplementary Table S10.

**Supplementary Table S10. Summary of two-way repeated measures ANOVA results for platelet calcium flux experiments.**

| Comparison | Effect Tested | F statistic | P value | Interpretation |
|---|---|---|---|---|
| **Calcium vs Calcium + aPC** | Time | F(120,360) = 4.376 | <0.0001 | Calcium signal changes over time |
| **Calcium vs Calcium + aPC** | Condition | F(1,3) = 0.1363 | 0.7365 | No effect of aPC above calcium alone |
| **Calcium vs Calcium + aPC** | Time × Condition | F(120,360) = 0.2765 | >0.9999 | No time-dependent difference between conditions |
| **0.1 U/mL Thrombin vs 0.1 U/mL Thrombin + FR900359** | Time | F(120,360) = 19.15 | <0.0001 | Time-dependent calcium response |
| **0.1 U/mL Thrombin vs 0.1 U/mL Thrombin + FR900359** | Condition | F(1,3) = 28.78 | 0.0127 | FR900359 significantly reduces response |
| **0.1 U/mL Thrombin vs 0.1 U/mL Thrombin + FR900359** | Time × Condition | F(120,360) = 18.81 | <0.0001 | FR900359 changes response over time |
| **0.5 U/mL Thrombin vs 0.5 U/mL Thrombin + FR900359** | Time | F(120,360) = 117.7 | <0.0001 | Time-dependent calcium response |
| **0.5 U/mL Thrombin vs 0.5 U/mL Thrombin + FR900359** | Condition | F(1,3) = 183.6 | 0.0009 | FR900359 strongly reduces response |
| **0.5 U/mL Thrombin vs 0.5 U/mL Thrombin + FR900359** | Time × Condition | F(120,360) = 127.5 | <0.0001 | FR900359 changes response over time |
